# Anatomically distinct cortical tracking of music and speech by slow (1-8Hz) and fast (70-120Hz) oscillatory activity

**DOI:** 10.1101/2024.06.12.598687

**Authors:** Sergio Osorio, M. Florencia Assaneo

## Abstract

Music and speech encode hierarchically organized structural complexity at the service of human expressiveness and communication. Previous research has shown that populations of neurons in auditory regions track the envelope of acoustic signals within the range of slow and fast oscillatory activity. However, the extent to which cortical tracking is influenced by the interplay between stimulus type, frequency band, and brain anatomy remains an open question. In this study, we reanalyzed intracranial recordings from thirty subjects implanted with electrocorticography (ECoG) grids in the left cerebral hemisphere, drawn from an existing open-access ECoG database. Participants passively watched a movie where visual scenes were accompanied by either music or speech stimuli. Cross-correlation between brain activity and the envelope of music and speech signals, along with density-based clustering analyses and linear mixed effect modeling, revealed both anatomically overlapping and functionally distinct mapping of the tracking effect as a function of stimulus type and frequency band. We observed widespread left-hemisphere tracking of music and speech signals in the Slow Frequency Band (SFB, band-passed filtered low-frequency signal between 1-8Hz), with near zero temporal lags. In contrast, cortical tracking in the High Frequency Band (HFB, envelope of the 70-120Hz band-passed filtered signal) was higher during speech perception, was more densely concentrated in classical language processing areas, and showed a frontal-to-temporal gradient in lag values that was not observed during perception of musical stimuli. Our results highlight a complex interaction between cortical region and frequency band that shapes temporal dynamics during processing of naturalistic music and speech signals.

## Introduction

Music and speech encode hierarchically organized structural complexity and support emotional and semantic expressiveness in human communication (1–7). To achieve this goal, small structural units (e.g., notes or syllables) are linearly combined to build up longer units (e.g., phrases or sentences), establishing non-linear relationships between high-level abstract representations. Hence, understanding how the brain physiologically supports perception of music and speech is of great importance for cognitive and auditory neuroscience.

Because both signals deploy in time, hierarchical structure is physically embedded in the modulation of a signal’s amplitude over time (i.e. the acoustic envelope), which can be quantitively assessed via power density and Fourier analyses. Across multiple languages and different musical genres, music and speech signals are characterized by spectral peaks at around 2Hz and 5Hz, respectively, which roughly correspond to the mean note rate (8,9) and the mean syllabic rate (9,10). Despite subtle differences in their temporal dynamics, perception of both music and speech requires the ability to track multiple structural units and their dependencies. This has led to propose that the neural architecture supporting perception of both signals must derive from similar evolutionary affordances and must be supported by shared neural mechanisms (2,11,12).

During the perception of music and speech, neuronal populations track temporal regularities across different hierarchical levels within the signal, a phenomenon known as cortical tracking. Cortical tracking of structural units, such as notes and phrases, and of acoustic features, such as rhythm, melody, and modulations in the signal’s envelope, has been observed within the Slow Frequency Band (SFB, band-passed filtered low-frequency signal between 1-8Hz) activity during perception of musical stimuli (13–17). This tracking has been shown to be modulated by several factors, including note rate (14), musical expertise (13), stimulus familiarity (18) and stimulus complexity (19). Similarly, during speech perception, brain activity within the same frequency band (i.e., SFB) tracks structural units, such as syllables, words and phrases, and acoustic features such as word onsets and amplitude modulations in the signal’s envelope (17,20–26). SFB tracking of speech signals is modulated by factors such as syllabic rate (20,22), speech intelligibility (22,27,28) and attention (29,30).

Neural tracking of music and speech has also been observed within the range of High Frequency gamma Band (HFB, >70 Hz) activity, particularly through the modulation of HFB amplitude (22,23,31–34). During perception of music, cortical tracking of the acoustic envelope and various other acoustic features, including pitch, voice, and melody, has been associated with both time-locked and sustained HFB neural responses in regions such as the superior temporal, middle temporal, supramarginal, and dorsolateral prefrontal cortices (32,33,35–38). Similarly, during speech perception, HFB tracking of structural units, such as syllables, words and phrases, and of the acoustic features such as pitch, peak rate, syllable onsets, and the signal’s envelope, has been observed in the superior temporal regions and ventrolateral prefrontal cortices (22,23,31,34,39,40), with a gradient of increasing tracking magnitude from perisylvian and prefrontal areas toward primary auditory regions (31). Despite their limited spatial resolution and signal-to-noise ratios compared to intracranial recordings, MEG and EEG studies corroborate HFB tracking of the speech envelope in temporal electrodes and source-reconstructed auditory areas (41,42).

Studies using invasive methods such as electrocorticography and stereotactic EEG have demonstrated that neural tracking of acoustic signals is widely distributed in cortical space, exhibiting little or no anatomical regional selectivity based on stimulus type (35,36,39). This findings contrasts with findings from fMRI studies, which indicate anatomical specialization for speech in ventrolateral prefrontal regions and for music in dorsolateral prefrontal and inferior parietal regions (43–48). Despite this apparent discrepancy, recent evidence points to selectivity depending on the type of information being tracked. One previous study reported that posterior portions of the Superior Temporal Gyrus (STG) are more sensitive to syntactic complexity in speech stimuli, whereas posterior portions of the Middle Temporal Gyrus (MTG) show greater sensitivity to music complexity (49). Furthermore, selectivity across frequency bands has also been observed: while the envelope of the music and speech acoustic signals is similarly tracked by both SFB and HFB activity, acoustic edges, which correspond to the peak rate of temporal modulations in the acoustic signals, are more accurately predicted by activity in the SFB (39). These findings therefore suggest a more intricate interplay between stimulus type, frequency band, and brain anatomy, highlighting the need for further investigation to fully characterize these interactions.

Here, we analyzed an open-access database of electrocortical recordings acquired during naturalistic, multimodal perception of music and speech stimuli (50). Prior to surgical procedures, patients passively watched 30-second videos containing either music (multi-instrument clips) or speech (single voice and dialogues) audios, while brain activity was recorded. Data from a subsample of 30 subjects with electrode grids located in left prefrontal, motor, premotor, somatosensory and/or temporal regions was analyzed, thus offering a coverage of most left-hemisphere cortical areas involved in music and speech perception. Specifically, we aimed to understand the anatomical organization and temporal dynamics underlying the neural tracking of music and speech, examining both slow (1-8Hz) and fast (70-120Hz) oscillatory bands known to be involved in perception and processing of both acoustic signals.

To quantify cortical tracking of music and speech in a straightforward way, we conducted cross-correlation analyses between the cochlear envelope of acoustic signals and the bandpass-filtered brain signals (SFB, the brain activity filtered between 1-8 Hz; and HFB the amplitude modulation of the 70-120 Hz filtered brain signal). To investigate whether frequency-band-dependent tracking effects can be mapped to similar or different cortical areas, we conducted density-based clustering analyses. Finally, to investigate the effect of condition and cortical region in the strength of the tracking effect and its temporal dynamics, we conducted mixed effect modeling analyzes using maximum correlation and temporal lags as the predicted variables, and frequency band (SFB, HFB), stimulus type (music, speech), and anatomical location of electrodes (cortical region) as regressors, while controlling for the within-subject nature of the data and differences in EcoG grid locations across subjects as potential random effects.

## Materials and methods

Data was obtained from a publicly available dataset (50) collected at the University Medical Center Utrecht from 63 patients that underwent intracortical electrode implantation as part of diagnosis procedures for resection surgery due to a clinical history of drug-resistant epileptic seizures. For this study, we used data from a subsample of 30 subjects (mean age = 27.33, SD=15.28, 19 females, see supplementary table s1), selected on the basis of electrode grid coverage (i.e., only patients with grids in left prefrontal, motor, premotor, somatosensory and/or temporal regions). Language dominance for all subjects was localized to the left hemisphere (table s1). This resulted in a total number of 1,858 electrodes across all subjects (mean = 61.93, SD = 19.09, supplementary table s2) distributed across left-hemisphere prefrontal, central and temporal regions (figure s1).

Participants were passively presented 30-second interleaved excerpts from the movie “Pipi on the run” (Pårymmen med Pippi Långstrump, 1970) dubbed to the Dutch language, where visual scenes were accompanied by music (7 blocks) or speech (6 blocks) stimuli. All visual and auditory excerpts were different from one another. The task was originally designed as part of a language mapping procedure prior to surgical intervention. All participants and legal guardians, when applicable, provided their informed consent to participate in the study and to make their de-identified data publicly available (50). Procedures were revised and approved by the Medical Ethical Committee of the University Medical Center Utrecht in accordance with the Declaration of Helsinki (2013).

## Data analysis

### Acoustic signals

Acoustic data was analyzed using MATLAB Version 9.6.0 (2019a). The acoustic signal was extracted from the audiovisual stimulus and segmented according to stimulus type (speech or music). The last musical segment was excluded from analyses to have a balanced number of segments across the two conditions. The segmented signals were then down-sampled to 16,000 Hz. Next, each segment’s auditory representation in the human cochlea was obtained by detrending, resampling (200hz), and filtering each signal in 128 logarithmically spaced frequencies within the range of human audition (180 to 7,246 Hz). The cochlear spectrogram was obtained using the NSL toolbox (51). Finally, signals were averaged across the 128 frequencies to obtain the cochlear envelope of each acoustic segment. For each envelope, the Power Spectrum Density was obtained using Welch’s method.

### EcoG and neuroanatomical data preprocessing

Electrophysiological and neuroanatomical data was preprocessed using Brainstorm (52). Electrodes were first re-referenced to the local average of each individual EcoG grid. Next, data was bandpass filtered between 0.5 and 120Hz using an FIR Keyser filter (attenuation level = 60 dB, Order = 3714). A second-order IIR notch filter was applied to remove line noise (50Hz) and harmonics (100Hz). Then, electrodes reported as bad in the original dataset and additional noisy channels visually identified on the basis of their power spectrum properties via Welch’s PSD analysis were rejected from further analysis (mean = 3.5, SD = 4.6).

To inspect EcoG grid location, electrodes were plotted against each subject’s cortical surfaces, which are available in the original dataset. After replicating individual electrode locations for each participant, we then projected all electrodes in a the MNI-ICBM152 cortical template (figure s1). In a few cases, this procedure resulted in spatial distortion of some electrodes due to conversion between coordinate systems. In such cases, electrode location was corrected by visually comparing the electrode’s position in the subject’s anatomy and relocating it if necessary to the corresponding anatomical area in the common cortical space. For more details on structural MRI preprocessing and post-surgical grid corregistration procedures in the original data, see Berezutskaya et al., 2022.

Electrophysiological signals were then bandpass-filtered and epoched using Fieldtrip (53). A third order Butterworth IIR filter was used to obtain continuous brain signals within the Slow Frequency Band (SFB, 1-8Hz) and High-Frequency Band (HFB, 70-120Hz). For the latter frequency band, the envelope was obtained by extracting the absolute value of the signal’s Hilbert transform. Finally, data was epoched into twelve 30-second segments, corresponding to the six music and six speech clips.

### Cross-correlation analysis

Cross-correlation analyses were conducted using MATLAB Version 9.6.0 (2019a) and 9.14.0 (2023a). The cross-correlation function was estimated for each electrode between the z-normalized brain signals and the z-normalized cochlear envelopes of the music and speech segments. This was done per subject, for the two frequency bands of interest (SFB and HFB). Previous studies have used small temporal windows of approximately 500ms, which excludes the slowest temporal modulations in the acoustic signals. Here, we used a maximum lag of ± 400 samples, corresponding to a four-second window between -2000 and 2000ms. This is in line with findings that larger temporal windows are associated with better model performance in decoding analyses of brain signals using Artificial Neural Networks (54). In the current work, no analyses were conducted between the brain signals and information in the visual modality.

The maximum correlation coefficient and its corresponding temporal lag were extracted from the correlation function. The mean coefficient estimates and corresponding temporal lags were obtained by averaging across the values for the six segments within each condition at the subject level (figure 1b, for data distributions see supplementary figure s2). Statistical significance was estimated for each individual by obtaining a null distribution of cross-correlation coefficients via permutation (figure 1b). For each subject, the cross-correlation function was estimated 1,000 times between the z-normalized brain signals and the z-normalized cochlear envelopes of six randomly generated white-noise signals per condition. P-values for the observed data were subsequently obtained by estimating the probability of observing the same or more extreme cross-correlation coefficients given the empirically-obtained null distributions. The threshold for statistical significance was set at an alpha value of 0.001.

**Figure 1.**
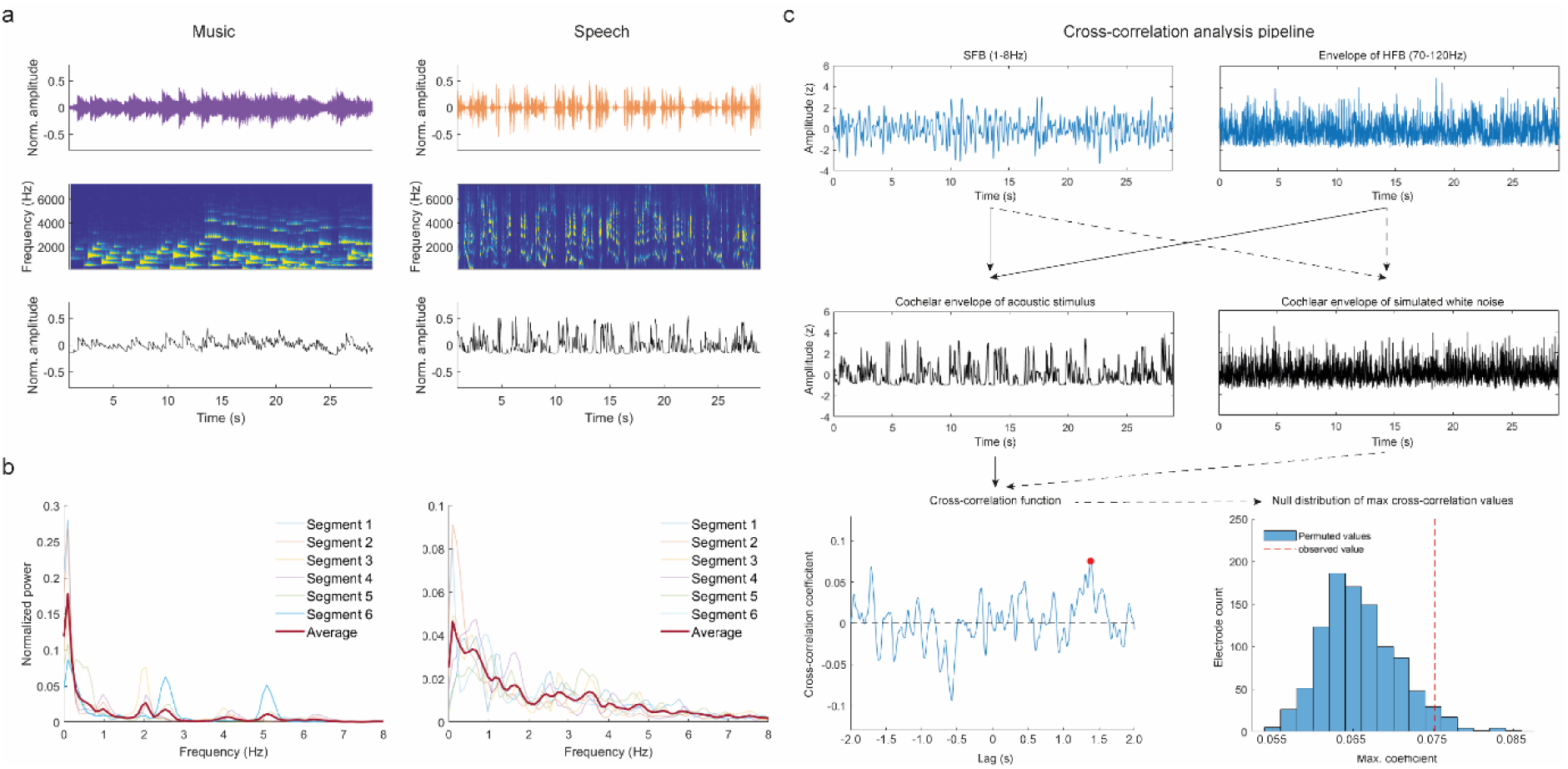
Analysis of acoustic signals. **a.** Sound waves of representative acoustic segments for music (top-left panel) and speech (top-right panel), their cochlear spectrograms (middle panels) and cochlear envelope (bottom panels). **b.** PSD for all music (left) and speech (right) segments. For normalization, power at each frequency bin was divided the sum of power values across all frequencies. The red line shows average power across segments. **c.** Schematic representation of the cross-correlation analysis. For brain signals (SFB, 1-8Hz, top left and envelope of HFB, 70-120Hz, top right), cross-correlation was estimated against the cochlear envelope of stimuli signals (middle left) to obtain the maximum correlation coefficient and its corresponding lag (bottom left). A permutation procedure was conducted by estimating the cross-correlation function between SFB and HFB brain signals and the cochlear envelope of simulated white-noise (middle right), to obtain a null distribution of maximum correlation coefficients which was used to estimate significance thresholds. Solid black arrows represent the pipeline for real data. Dotted black lines represent the permutation procedure.

### Density-based cluster analysis

All statistically significant electrodes across our 30 subjects were then plotted in a normalized cortical space (MNI-ICBM152). Electrodes that significantly correlated with the acoustic envelope were identified performing multiple comparisons (i.e., 1858 electrodes were studied). Accordingly, a high number of false positives are expected, even if a more conservative alpha value was chosen a priori. Because the permutation procedure is agnostic of the anatomical location of electrodes, and in order to identify cortical regions where the group-level effect was localized, we additionally conducted a density-based cluster classification analysis on the collapsed data across all subjects using the DBSCAN algorithm (55) as implemented by MATLAB. The assumption was that the overall effect of cortical tracking should be more pronounced in task-relevant cortical regions. DBSCAN should therefore spatially constrain any statistical effect in a data-driven manner to functionally relevant cortical areas, while taking care of isolated electrodes which would otherwise be considered outliers.

DBSCAN requires only two initial parameters: the minimum number of data points for a cluster to be identified and the minimum radius within which those data points should be contained (i.e., the epsilon parameter). This means that no starting data point or number of clusters need to be defined a priori, unlike other clustering methods. To derive the optimal parameters for the DBSCAN algorithm, we conducted a procedure to obtain the parameter combination that optimized clustering within each condition. We tested five minimum points (12 to 20 in steps of 2) and a range of 21 epsilon parameters (0 to 0.04 in steps of 0.002). A matrix of parameter optimization indices (*OI*) was obtained by the following formula.

Where Є is the 1-by-21 vector containing the list of possible minimum radiuses, ***P*** is the 1-by-5 vector of possible minimum data points, Where ***E*** is the 5-by-21 matrix containing the percentage of electrodes obtained for each combination of initial parameters, and ***C*** is the 5-by-21 matrix containing the number of clusters obtained by each combination of parameters. Finally, we normalized ***OI*** so that values range between 0 and 1, with 0 indicating poor algorithm performance and 1 indicating optimal performance.

Supplementary figure s3 shows the normalized OI values as a function of different parameter combinations for the SFB and HFB analyses. The ideal parameters are those which maximize *OI_norm_* with the lowest possible cluster radius and the highest possible number of electrodes within such cluster. The final parameters used for DBSCAN clustering analyses for each frequency band and stimulus type are provided in table 1.

**Table 1.**
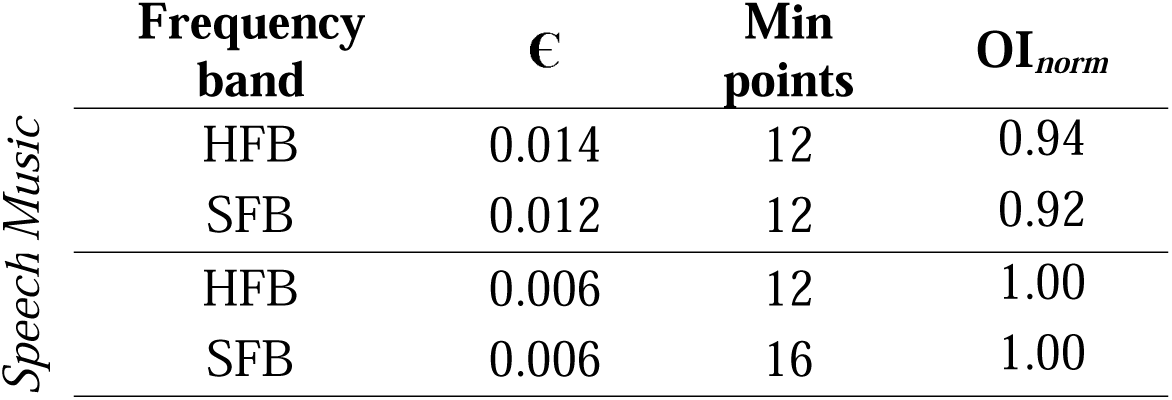
Optimal parameters for density-based clustering analysis. The epsilon (Є) parameter represents the radius of search, and the minimum points stands for the minimum number of electrodes within the search area for cluster identification.

### Electrode anatomical labeling

DBSCAN returns a numbered list of identified clusters under the specified parameter combination. We therefore grouped all clusters resulting from DBSCAN analyses as a single group and anatomically labeled each electrode using the Desikan-Killiany anatomical atlas (56). We took this approach because clusters where the result of an algorithmic operation performed in space that is agnostic of brain anatomy, which in practice means that two small clusters could be identified within the same anatomical region. Electrodes were therefore labeled according to the cortical area in which they were located by plotting each electrode on the MNI-ICBM152 template against overlaid parcellations from the Desikan-Killiany atlas. This resulted in nine cortical regions of interest (table 2), which were used as categorical predictors in our linear mixed effect modeling analyses.

**Table 2.**
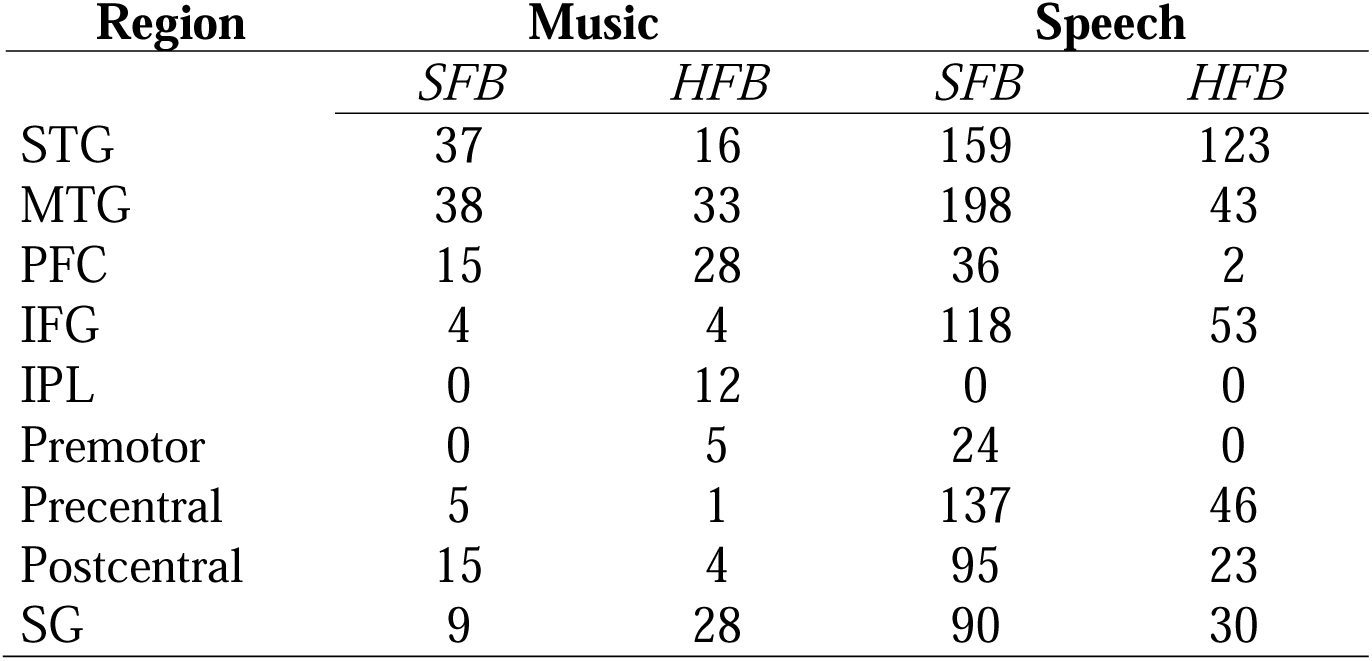
Number of statistically significant electrodes per anatomical region. *STG*, (Superior Temporal Gyrus), *MTG* (Middle Temporal Gyrus), PFC (Prefrontal Cortex) *IFG* (Inferior Frontal Gyrus), *IPL* (Inferior Parietal Lobe), SG (Supramarginal Gyrus)

| Region | Music |  | Speech |  |
| --- | --- | --- | --- | --- |
|  | *SFB* | *HFB* | *SFB* | *HFB* |
| STG | 37 | 16 | 159 | 123 |
| MTG | 38 | 33 | 198 | 43 |
| PFC | 15 | 28 | 36 | 2 |
| IFG | 4 | 4 | 118 | 53 |
| IPL | 0 | 12 | 0 | 0 |
| Premotor | 0 | 5 | 24 | 0 |
| Precentral | 5 | 1 | 137 | 46 |
| Postcentral | 15 | 4 | 95 | 23 |
| SG | 9 | 28 | 90 | 30 |

### Linear mixed effect modeling

Linear Mixed Effect (LME) modeling was conducted using the LME4 package (57) in R (R Core Team, 2023) using R Studio version 2023.6.1 (R Studio Corte Team, 2023). Data was modeled for the two response variables of interest: cross-correlation values and temporal lags. For each LME analysis, the minimum number of datapoints within each cortical region for inclusion in the model was set to nine. For music, electrodes within the STG, MTG, PFC and Supramarginal Gyrus (SG) were included in the analysis. The IFG, premotor cortex, precentral and postcentral gyri were excluded because not enough electrodes were found within these regions across frequency bands. For speech, electrodes within the STG, MTG, precentral and postcentral gyri, IFG, and SG were included in the analysis. Not enough electrodes were found across frequency bands in the premotor or prefrontal cortices, so these regions were excluded. We additionally performed a joint LME analysis for music and speech including the STG, MTG and SG as cortical regions of interest.

A forward stepwise model selection procedure using Maximum Likelihood Estimation was conducted to identify the models that best fit our data (table 3 and supplementary tables s3, s4 and s5). Single predictor models for frequency band and cortical region including a random intercept for subject were tested against a null (random intercept-only) model. If any single-predictor model was significantly better than the null model, the effect of adding additional terms and interactions while keeping the random intercept unchanged was tested. Once the optimal combination of fixed effect terms was determined, the effect of adding other random effects was tested. This procedure was conducted for music and speech separately. However, we additionally conducted a joint LME analysis where condition (music and speech) was considered an additional fixed effect. Table 3 shows the best models predicting cross-correlation coefficients and temporal lags per condition with their corresponding Akaike Information Criterion (AIC) and Bayes Information Criterion (BIC). For music, no model outperformed a null model in predicting temporal lags. For more information on model selection procedures, see supplementary materials.

**Table 3.** Best fit models for each condition and for the collapsed model and for each response variable of interest. For music, no model was significantly better in predicting temporal lags than a null model.

|  | LME model | AIC | BIC |
| --- | --- | --- | --- |
| Music | r ~ frequency + (1\|subject) | -1299,136 | -1285,854 |
|  | lag ~ (1+subject) | 277.78 | 287.73 |
| Speech | r ~ frequency*region + (1\|subject) + (1\|subject:region) | -6351,467 | -6276,218 |
|  | lag ~ frequency*region + (1\|subject) + (1\|subject:region) | 13426,74 | 13501,99 |
| Both | r ~ frequency*region*condition + (1\|subject) + (1\|subject:region) | -4379.39 | -4309.04 |
|  | lag ~ frequency*region*condition + 1(subject) | 9624.23 | 9689.89 |

## Results

We first studied the properties of our acoustic stimuli. Top panels in figure 1a show the soundwave of two representative music and speech acoustic segments (see also figure s4 in supplementary materials for all other segments). A cochlear filter was applied to obtain the approximate spectrotemporal representation of each music and speech segment in the human cochlea (figure 1a, middle panels, see methods). Next, the cochlear envelope of each segment was estimated by averaging across all the cochlear frequencies (figure 1a, bottom panels). Figure 1b shows the normalized spectral power for the six music (left) and speech (right) segments correspondingly, obtained via Welch’s Power Spectrum Density (PSD) method. PSD analyses show high variability in spectral peaks across the different music and speech segments, consistent with the uncontrolled nature of the stimuli. The mean spectrum across music cochlear envelopes, however, shows peaks at around 0.2, 2 and 5 Hz, suggesting that across segments, music signals are somewhat more rhythmic than speech signals. Analyses confirmed statistically significant weak-to-moderate positive correlations between the six acoustic envelopes of music (α = 0.05, two-tailed, table 4, top panel). For speech, no clear dominant peak was observed when the cochlear envelopes were averaged, reflecting marked differences in the temporal structure across the six different speech segments. This is consistent with the nature of the speech stimuli, which included both monologues and conversations between multiple individuals. Correlation analysis showed weak negative correlations for the acoustic envelopes of speech (table 4, bottom panel).

**Table 4.** Correlation across the music and speech segments.

|  |  |  |  |  |  |
| --- | --- | --- | --- | --- | --- |
| 2 | 0.397*** |  |  |  |  |
| 3 | 0.232*** | 0.103*** |  |  |  |
| 4 | 0.442*** | 0.560*** | 0.299*** |  |  |
| 5 | 0.290*** | 0.253*** | 0.012_n.s. | 0.225*** |  |
| 6 | 0.029*** | -0.131*** | 0.147*** | -0.037*** | -0.006_n.s. |
| 1 |  |  |  |  |  |
| 2 | 0.036_n.s. |  |  |  |  |
| 3 | 0.030*** | 0.006_n.s. |  |  |  |
| 4 | -0.022_n.s. | 0.030*** | -0.050*** |  |  |
| 5 | 0.049*** | -0.102*** | -0.042*** | -0.098*** |  |
| 6 | 0.043*** | 0.164*** | -0.059*** | 0.002_n.s. | -0.006_n.s. |

Speech
\*\*\* $p < 0.001$ , \*\* $p < 0.01$ , \* $p < 0.05$

We then studied neural tracking of the music and speech envelopes. For this, we estimated the cross-correlation function between the brain and acoustic signals. Figure 1c shows a schematic representation of the cross-correlation analysis pipeline. For each subject (n = 30), we obtained the cross-correlation function between the cochlear envelopes of the music (n = 6) and speech (n = 6) segments, and the brain signal recorded at each electrode (n = 1858, mean = 61.93, SD = 19.09) in the Slow Frequency Band (SFB, 1 to 8Hz bandpass filtered signal) and the High Frequency Band (HFB, envelope of the 70 to 120Hz bandpass filtered signal). For each cross-correlation, we then extracted the maximum brain-to-stimulus coefficient and its corresponding temporal lag. We finally averaged the coefficients across the six segments within each stimulus type. Statistical significance was estimated using a null distribution of cross-correlation coefficients obtained via permutation (n _permutations_ = 1000, α = 0.001, figure 1c). Finally, we additionally implemented a density-based clustering classification analysis using the DBSCAN algorithm to remove spatial outliers and identify cortical regions where significant electrodes were most densely grouped.

In both frequency bands, permutation and density-based clustering analyses showed a higher number of statistically significant electrodes tracking speech compared to music (figure 2). Electrodes tracking the music envelope in the SFB concentrated in perisylvian, temporal and prefrontal regions, including the middle and posterior portions of the STG, anterior, middle, and posterior portions of the MTG and ITG, the lowermost portions of the precentral and postcentral gyri, and the most anterior portion of the dorsolateral Prefrontal Cortex (dlPFC, figure 2a, left). For speech, cortical tracking in the SFB was abundantly found in perisylvian, ventral prefrontal, pre and postcentral, and temporal regions, including the anterior, middle, and posterior portions of both the STG and MTG, the lowermost portions of the SG, middle and lower portions of the precentral and postcentral gyri, premotor areas, the IFG, and the dlPFC (figure 2a, middle). In the SFB, the number of statistically significant electrodes tracking the speech signals (n = 823, percent from total = 44.22%) was 6.5 times higher than the number of electrodes tracking the music signals (n = 125, percent from total = 6.72%, figure 2b). Interestingly, within the SFB, 84% of the electrodes tracking the music signals (n = 105, percent from total = 5.65%) also tracked the envelope of the speech signals (figure 2a, right and figure 2b).

**Figure 2.**
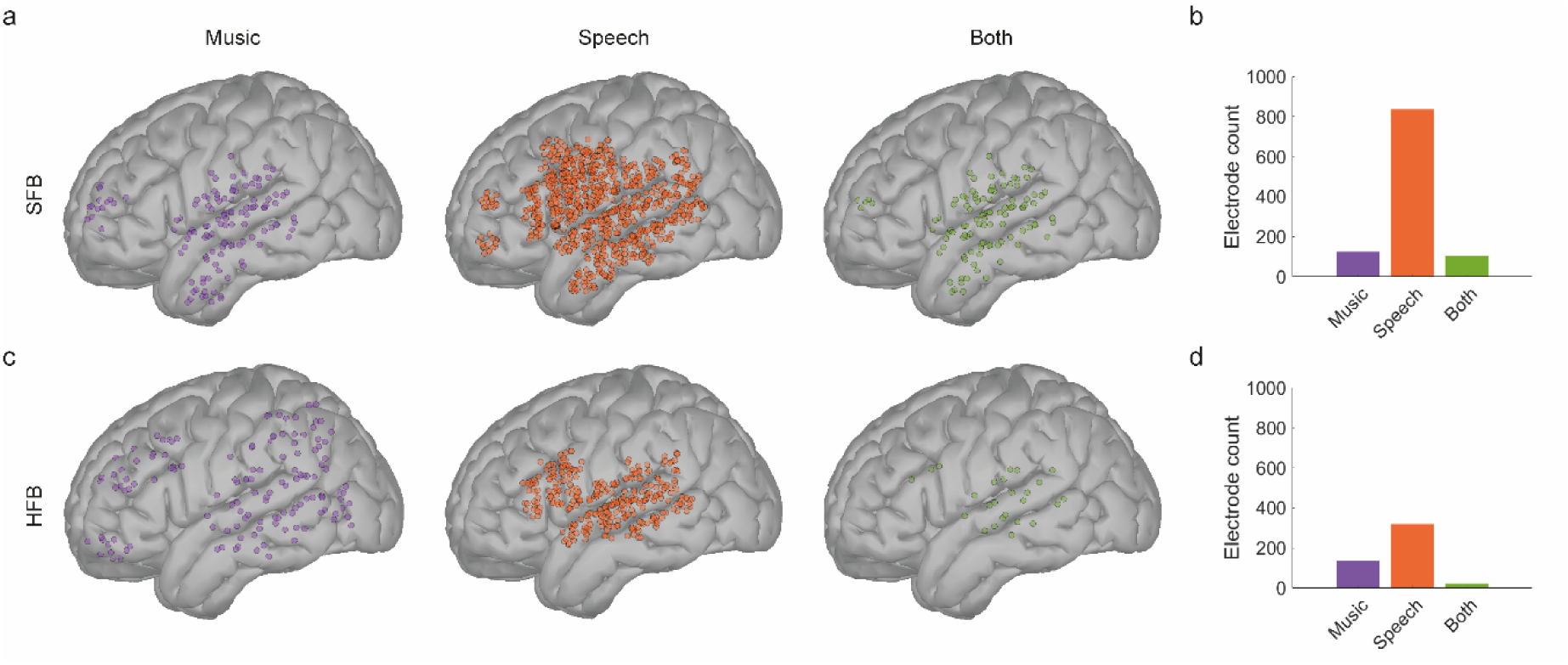
Statistically significant electrodes and their anatomical localization in the MNI-ICBM152 cortical template. **a.** and **c.** Spatial distribution of significant electrodes in the SFB (**a**) and HFB (**c**) after density-based clustering analyses for music (orange) and speech (purple). Electrodes that show mixed-selectivity (i.e., respond to both stimuli) are presented in green. **b.** and **d.** Number of electrodes that survive permutation statistics and density-based clustering classification in the SFB (**b**) and HFB (**d**).

Electrodes tracking music stimuli in the HFB concentrated around PFC and premotor areas, as well as in SG, IPL, STG and MTG regions (figure 2c, left). In turn, electrodes tracking speech in the HFB range were more densely concentrated in perisylvian regions, including the middle and posterior STG, lowermost portions of the precentral and postcentral gyri, and the middle and posterior portions of the IFG (figure 2c, middle). There was an approximate two-fold increase in the number of electrodes showing statistically significant tracking of speech in the HFB (n = 320, percent from total = 17.22%) compared to music (n = 136, percent from total = 7.32%, figure 2d). Only 18.32% of electrodes tracking the music envelope also tracked the speech envelope in the HFB. These electrodes were restricted to posterior portions of the STG (n = 25, percent from total = 1.34%, figure 2c, right). For a non-thresholded figure showing the raw distribution of cross-correlation coefficients and temporal lags prior to statistical analyses, see supplementary figure s5.

Next, we investigated the organization of cross-correlation values and temporal lags (figure 3). For tracking of music signals in the SFB, cross-correlation coefficients (mean = 0.09, SD = 0.006) showed a gradient of increasing values from association areas, including anterior portions of the ITG and MTG, towards primary auditory areas and posterior portions of the STG (figure 3a, top-left). Temporal lags were maximally concentrated around time zero (mean = 12ms, median = 50ms, SD = 470ms, figure 3a, top-right), following a similar gradient with more positive lags found in anterior and middle portions of the STG (figure 3c, left). In the HFB, coefficients (mean = 0. 096, SD = 0.013) showed a more spatially-distributed gradient, with increasing values towards association areas, including superior parietal, middle temporal, and prefrontal regions (figure 3a, bottom-left). Similar to the SFB, temporal lags for the HFB concentrated around time zero (mean = -14ms, median = 57ms, SD = 480ms, figure 3a, bottom-right), with maximum lags in middle and posterior portions of the STG (figure 3c, right).

**Figure 3.**
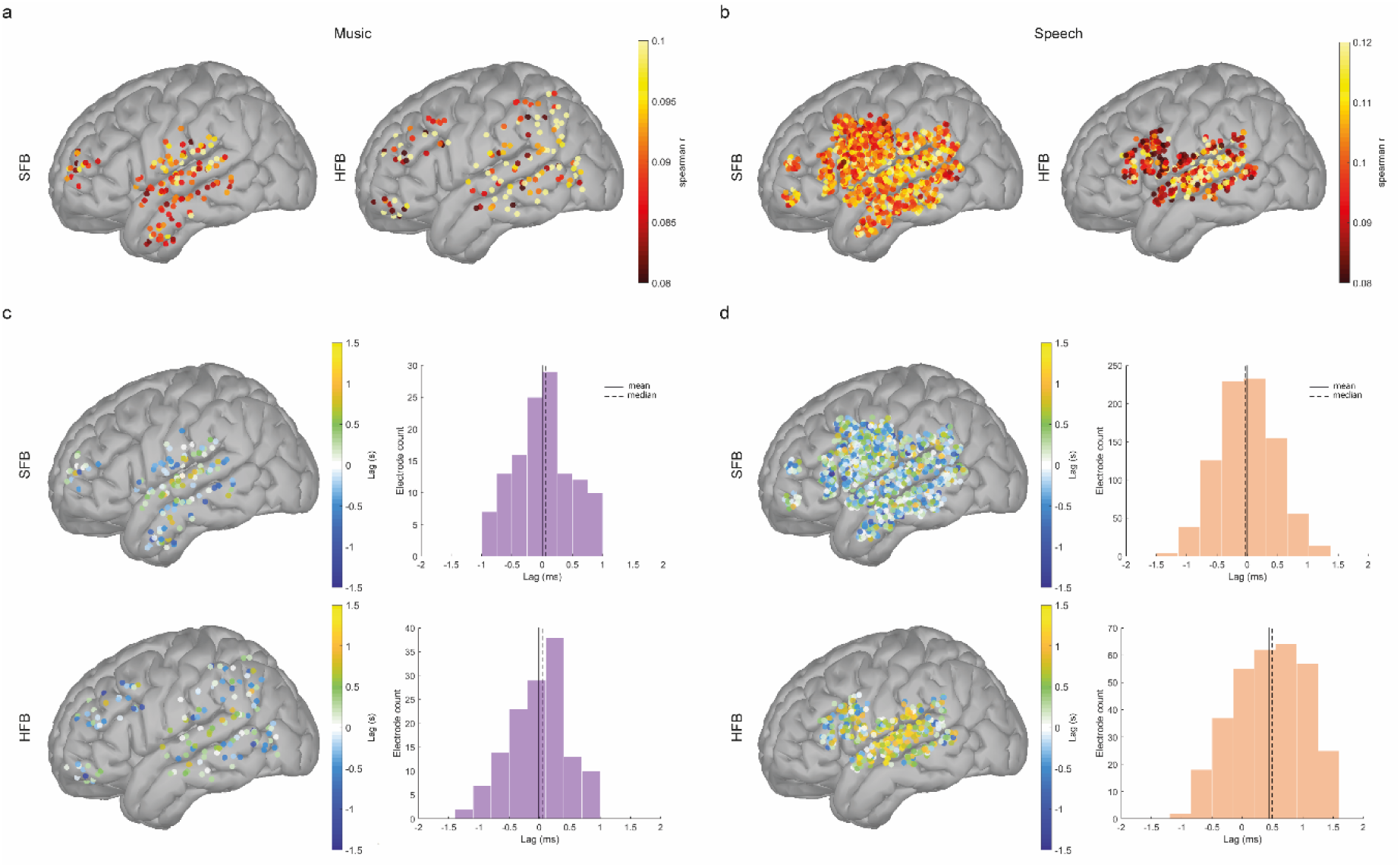
Cortical tracking of music and speech envelopes in the MNI-ICBM152 cortical template. **a.** and **b.** Spatial distribution of mean correlation values for music (a) and speech (b) in the SFB and HFB. **c.** and **d.** Spatial distribution and histograms (left) of temporal lags (right) in the SFB and HFB. Positive lags indicate that the acoustic signal precedes brain signals, whereas negative lags indicate that the brain signals precede the acoustic signal. All electrodes shown are statistically significant (p < 0.001, uncorrected) and survive clustering analyses.

The cortical tracking effect for the speech signal in the SFB (mean = 0.10, SD = 0.001) was particularly distributed in cortical space, with higher coefficients found along the anterior-to-posterior axis of the STG and MTG, but also in portions of the IFG, precentral gyrus and SG (figure 3b, top-left). Similar to music, temporal lags for speech in the SFB band converged around zero (mean = -6ms, median = -29ms, SD = 470ms, figure 3b, top-right, figure 3d, left). In turn, cortical tracking of the speech signal in the HFB (mean = 0.10, SD = 0.023) showed a densely-grouped distribution and sharp increase in cross-correlation coefficient values from prefrontal to posterior temporal regions, with a concentration of the highest cross-correlation coefficients in the middle and posterior STG (figure 3b, bottom-left). Interestingly, unlike all other conditions, mean temporal lags in the HFB band were biased towards positive values (mean = 440ms, median = 490ms, SD = 590ms, figure 3b, bottom-right). Temporal lags in the HFB followed a similar gradient as cross-correlation coefficients, with more positive lags (i.e. higher delays) concentrating around the middle portion of the STG (figure 3d, right). Finally, to investigate the consistency of our results across stimuli type, we re-analyzed our data using the same analytical pipeline, while randomly excluding two segments (corresponding to 33% of the data) at each of ten iterations. Results of these analyses show that both the magnitude of the effect as well as its spatial organization remain consistent within each stimulus type across all iterations (figure s6, supplementary materials).

To quantify the effect of cortical tracking more accurately across anatomical regions, electrodes were next labeled according to their anatomical location (see methods). Electrodes were classified within one of nine left-hemisphere cortical areas (figure 4a and table 2). Cortical tracking of music was more prominent within the MTG and STG regions in the SFB, and within the MTG, SG and PFC in the HFB (figure 4b). For speech stimuli, cortical tracking was higher within MTG, STG and postcentral regions for the SFB, and within STG and IFG for the HFB (figure 4c). Next, we resorted to mixed effect modeling to investigate how anatomical location and frequency band influence cortical tracking. The two response variables of interest were cross-correlation coefficients and temporal lags. The fixed effects of interest were frequency band and cortical region. The random effects of interest were subject (to account for the within subject nature of the task) and the interaction between subject and cortical region (to account for the different electrode grid locations across subjects). We performed a forward stepwise model selection procedure to select the best model explaining the data (see supplementary materials).

**Figure 4.**
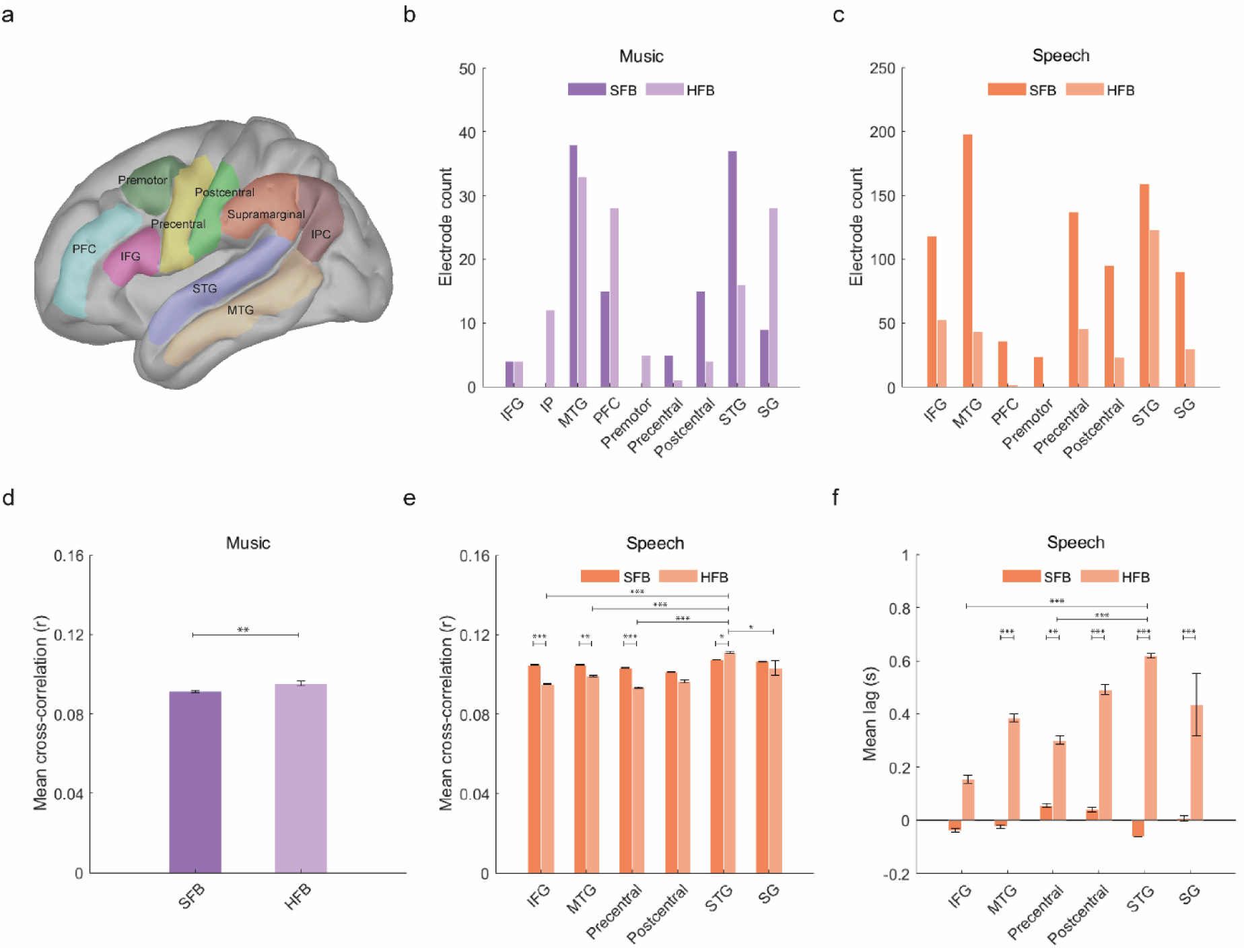
Anatomical regions of interest and significant effects as per mixed effect modeling analyses. **a.** Anatomical location of statistically significant electrodes after density-based clustering analyses. **b.** and **c.** Number of electrodes per anatomical parcellation for music (**a**) and speech (**c**). **d.** Main effect of frequency band for mean cross-correlation values during cortical tracking of music. **e.** Main and interaction effects for cross-correlation values during cortical tracking of speech. **f.** Main and interaction effects for temporal lags during cortical tracking of speech. Whiskers represent the Standard Error of the Mean (SEM). For all panels, * p < 0.05, ** p < 0.01, *** p < 0.001.

For music, a model including frequency band as fixed effect and subject as random effect best fit the observed cross-correlation coefficients (AIC = -1299.13, BIC = -1285.85, R^2^ = 0.12, R^2^ = 0.04, table s3). This model showed a statistically significant main effect of frequency (F (1, 184.87) = 9.00, p = 0.003, figure 4d). Estimated Marginal Means (emmeans) suggested that cortical tracking of the acoustic signals was significantly higher in the HFB (emmean = 0.094, C.I. = [0.092, 0.097]) than in the SFB (emmean = 0.089, C.I. = [0.086, 0.092]). However, no model was significantly better than a null model in predicting temporal lags during cortical tracking of music.

For speech, a model considering the interaction between frequency band and cortical region as fixed effect, and both subject and the subject-by-region interaction as random effects best explained the cross-correlation coefficients (AIC = -6351.47, BIC = - 6276.22, R^2^ = 0.19, R^2^ = 0.08, table s4). This model showed a significant main effect of frequency (F (1, 1066.49) = 25.73, p = 4.64e-07) and cortical region (F (5, 162.87) = 8.81, p = 2.01e-07), and a significant interaction between frequency and cortical region (F (5, 1058.98) = 7.47, p = 6.71e-07). Post-hoc pairwise comparisons showed that cortical tracking of the speech envelope was significantly higher in the SFB compared to the HFB within the IFG (emmean _SFB_ = 0.105, CI = [0.102, 0.108]; emmean _HFB_ = 0.095, CI =[0.091, 0.099], p < 0.0001), MTG (emmean _SFB_ = 0.10, CI = [0.101, 0.107]; emmean _HFB_

= 0.99, CI = [0.091, 0.010], p = 0.022) and precentral gyrus (emmean _SFB_ = 0.104, CI = [0.101, 0108]; emmean _HFB_ = 0.0942, CI = [0.0897, 0.099], p < 0.0001, figure 4e), and in the HFB compared to the SFB within the STG (emmean _SFB_ = 0.107, CI = [0.105, 0.110]; emmean _HFB_ = 0.102, CI = [0.108, 0.114], p = 0.016, figure 4e). In the HFB, cortical tracking was higher within STG (emmean = 0.109, CI = [0.107, 0.112]) compared to the IFG (emmean = 0.100, CI = [0.097, 0.103], p < 0.0001), MTG (emmean = 0.102, CI = [0.099, 0.105], p = 0.0002), precentral gyrus (emmean = 0.993, CI = [0.096, 0.102], p < 0.0001), postcentral gyrus (emmean = 0.993, CI = [0.095, 0.103], p = 0.0002) and SG (emmean = 0.104, CI = [0.100, 0.108], p = 0.040, figure 4e). No statistically significant differences were observed in cross-correlation coefficients across the different anatomical regions for the SFB.

For temporal lags during tracking of speech (figure 4f), the best model also included frequency, region and their interaction as fixed effects, and subject as well as the subject-by-region interaction as random effects (AIC = 1611.49, BIC = 1686.74, R^2^_Conditional_ = 0.25, R^2^_Marginal_ = 0.17, table s4). This model showed significant main effects of frequency (F (1, 1066.70) = 117.27, p < 2.20e-16), cortical region (F (5, 150.79) = 3.54, p = 0.0047) and the interaction between frequency and region (F (5, 1060.53) = 7.11, p = 1.48e-06). Post-hoc pairwise comparisons showed more positive lags for HFB compared to SFB within the IFG (emmean _HFB_ = 114ms, CI = [-33ms, 262ms]; emmean _SFB_ = -42ms, CI = [-152ms, 67ms], p = 0.050), precentral gyrus (emmean _HFB_ = 260ms, CI = [99ms, 420ms]; emmean _SFB_ = 18ms, CI = [9ms, 128ms], p = 0.004), MTG (emmean _HFB_ = 380ms, CI = [220ms, 541ms]; emmean _SFB_ = -23ms, CI = [-118ms, 72ms], p < 0.0001), postcentral gyrus (emmean _HFB_ = 431ms, CI = [219ms, 643ms]; emmean _SFB_ = 2ms, CI = [-122ms, 126ms], p = 0.0002), SG (emmean _HFB_ = 455ms, CI = [266ms, 643ms]; emmean _SFB_ = 32ms, CI [-9ms, 154ms], p < 0.0001) and STG (emmean _HFB_ = 618ms, CI = [510ms, 725ms]; emmean _SFB_ = -59ms, CI = [-158ms, 40ms], p < 0.0001), suggesting that delays between the acoustic envelope of speech and brain signals are higher in the HFB across all cortical regions of interest. In the HFB, tracking within the STG (emmean = 618ms) occurred 503ms later compared to the IFG (emmean = 114ms, p < 0.0001) and 358ms later compared to the postcentral gyrus (emmean = 260ms, CI = [100ms, 420ms], p = 0.002). No differences in temporal lags across cortical regions were observed within the SFB range (figure 4f).

Finally, we investigated the overall effect of cortical tracking and temporal lags by designing a joint model for music and speech. For this, we used a subsample of electrodes corresponding to the cortical regions where a significant effect of cortical tracking was observed for both music and speech. This included the STG, MTG and the SG (figure 5a). For mean cross-correlation values, the model that best fit the data was one including the interaction between condition, frequency band, and cortical region as fixed effects, and subject and the interaction between subject and cortical region as random effects (AIC = -4502.26, BIC = -4431.91, R^2^_Conditional_ = 0.26, R^2^_Marginal_ = 0.17, table s5). Results of this model suggest a main effect of condition (F (1, 799.99) = 76.49, p = 2.0e-16, figure 5b), where cortical tracking of speech (emmean = 0.105, C.I.= [0.103, 0.107]) is significantly higher than cortical tracking of music (emmean = 0.092, C.I. = [0.089, 0.095]). No effects were found for frequency band or cortical region. For mean temporal lags, the model included the interaction between condition, frequency band and region as fixed effects, and a random intercept for subject (AIC = 1191.96, BIC = 1257.61, R^2^ _Conditional_ = 0.22, R^2^ _Marginal_ = 0.20, table s5). Results of this model (figure 5c) show statistically significant main effects of frequency band (F (1, 787.56) = 14.16, p = 0.0002), condition (F (1, 776.45) = 13.27, p = 0.0003), and the interaction between frequency and condition (F (1, 749.70) = 35.98, p = 3.096e-09). Post-hoc pairwise comparisons showed that cortical tracking of speech in the HFB (emmean = 484ms, CI = [394ms, 574ms]) occurs, on average, 510ms later than tracking of speech in the SFB (emmean = -25ms, CI = [-85ms, 34ms], p < 0.0001), 504ms later than tracking of music in the HFB (emmean = -20ms, CI = [-147ms, 107ms], p < 0.0001), and 385ms later than tracking of music in the SFB (emmean = 99ms, CI = [-39ms, 237ms], p < 0.0001). No differences in temporal lags were observed for cortical tracking of music across frequency bands.

**Figure 5.**
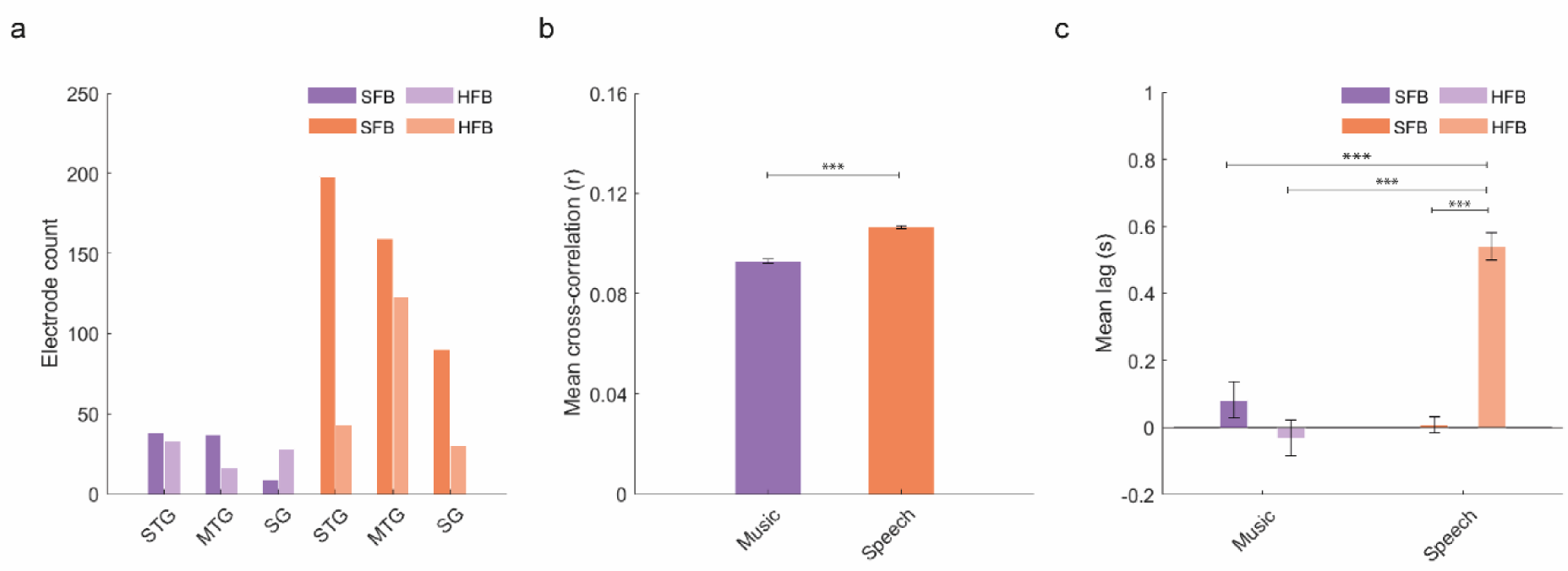
Number of electrodes per anatomical location and statistical effects of joint mixed effect model. Purple bars represent an effect for music whereas orange lines represent an effect for speech. **a.** Number of electrodes in the three anatomical location where statistically significant electrodes were found for both conditions. **b.** Main effect of condition for cross-correlation coefficients. **c.** Main effect and interaction for temporal lags. Whiskers represent the SEM. For all panels, * p < 0.05, ** p < 0.01, *** p < 0.001.

## Discussion

Cortical tracking of music and speech has been previously observed in the range of slow (1-8Hz, SFB) and fast (>70Hz, HFB) oscillatory activity under carefully controlled conditions (13–16,18–20,22,28,29) and under naturalistic perceptual scenarios (17,21,38,39,41,42,58–63,23,25,26,30,31,33–35). Here, we asked whether complex interactions between anatomical region, frequency band and stimulus type exist during perception of naturalistic music and speech stimuli that might have not been previously observed due to sample size, methodological or analytical limitations. For this, we leverage the high spatiotemporal resolution of electrocorticography by reanalyzing a publicly available EcoG dataset collected during passive perception of naturalistic music and speech stimuli. Our findings highlight both shared and distinct neural patterns between music and speech processing across different cortical regions, with marked differences in the spectral and temporal dynamics of brain responses.

### Acoustic Features of Music and Speech

The spectral power analysis of the cochlear envelopes revealed substantial variability across the six acoustic segments of music and speech. This variability was expected given the naturalistic nature of the stimuli. Despite this variability, the spectral power distributions of the cochlear envelopes suggest that music tends to be more rhythmic, as evidenced by the dominant peaks at approximately 0.2, 2, and 5 Hz in the music envelopes. In contrast, the speech segments did not exhibit a prominent peak, indicating more complex and variable temporal patterns, in line with the diverse types of speech stimuli employed in the study. Correlation analyses showed weak-to-moderate positive correlations within music envelopes, suggesting some degree of temporal coherence across music segments. This is further corroborated by the results showing weak negative correlations within speech envelopes, which reflects the highly variable temporal structures inherent to speech, including shifts in rhythm due to conversational interactions or changes in speech style.

### Cortical organization of the tracking effect in the slow and fast frequency bands

In line with similar intracortical studies (22,23,35,39), we found distributed cortical tracking in left hemisphere regions classically involved in language and music perception. Density-based clustering analyses showed overlapping tracking of music and speech in the SFB and HFB within the MTG, STG, SG, precentral and postcentral gyri. These findings are in line with previous research using source-localized MEG signals (20,21,62,64) and intracortical data (16,31–33,35,39). Overlap of cortical tracking around perisylvian and middle and superior temporal areas for both stimuli suggests that shared neuroanatomical mechanisms are involved in the processing of the acoustic envelopes of music and speech (2,11,12). This was particularly true for the SFB, where 84% of electrodes that showed a statistically significant effect for music also showed a significant effect for speech, thus suggesting not only high anatomical overlap, but also sensitivity to both types of stimuli within the range of slow oscillations.

However, we also found differences that were both frequency- and stimulus-dependent. Tracking of music in both the SFB and HFB, and of speech in the SFB, extended beyond classic auditory regions to include prefrontal, middle temporal, and parietal areas. In contrast, tracking of speech in the HFB was densely localized to perisylvian and ventral prefrontal regions, indicating a more focused cortical engagement for speech processing in the range of high gamma band activity. One previous EcoG study found similar a spatial organization pattern of neural activations during speech perception, with SFB activity showing a global distribution in cortical space, while HFB activity was primarily localized to the STG (23). This frequency-dependent pattern of global versus local spatial organization aligns with the idea that slow oscillatory activity facilitates communication among globally distributed neuronal ensembles, whereas high frequency activity reflects local activation of small neural populations in response to specific sensory features (65–73). Notably, the importance of cross-frequency interactions between the SFB and gamma band oscillations for speech perception and language processing has also been well documented (74–77). These interactions likely support multiscale extraction of both acoustic features and hierarchical structures that are essential for language comprehension. However, such cross-frequency coupling has been primarily observed between the slow oscillatory range (0.5–8 Hz) and the slower component of gamma-band activity (25–40 Hz), rather than the faster component (>70 Hz).

### Functional anatomical selectivity during tracking of natural music and speech signals

In the SFB, electrodes tracking the envelope of speech were observed within the IFG, which were nevertheless absent during tracking of music stimuli. In contrast, in the HFB, electrodes tracking the envelope of music stimuli were observed within the dlPFC and IPL regions, which were absent during tracking of speech. Both of these findings are in line with the fMRI literature on the role of the dlPFC and parietal cortex during music perception, and of the IFG during speech perception (43–48). In the EcoG literature, selective dlPFC recruitment in response to music has been previously reported during perception of musical stimuli, which has been attributed to attentional and emotional processing, as well as recall of complex melodic features in music stimuli (32,33,78). The involvement of the IFG during tracking of speech stimuli in both SFB and HFB has also been extensively reported in previous studies using intracortical methods (23,54,78). This suggests that in spite of partial anatomical overlap in perisylvian regions, cortical tracking of music and speech signals can also be intracortically mapped to distinct parietal and prefrontal brain regions in a stimulus-dependent manner.

Multiple roles have been previously attributed to the IFG during speech perception, including syntactic processing, semantic processing and phonological working memory (45,79–81). Admittedly, activation of the IFG has also been reported during music perception, however, recent metanalyses and clinical studies suggest this effect is right lateralized (82,83). Finally, increased BOLD activation of IPL regions has been observed in response to music compared to speech (84), and increased musical training is associated with increased IPL activity among musicians (85).

Not many EcoG studies have directly investigated the extent to which tracking of music and speech can be mapped to distinct cortical areas and, more importantly, how interactions between cortical regions, frequency bands and stimulus type influence this mapping. A recent study reported no substantial differences in anatomical regional selectivity in how brain signals track music and speech across delta (1-4Hz), theta (5-8Hz), alpha (8–12), beta (18-30Hz), low gamma (30-50Hz) and high-gamma (80-150Hz) frequency bands (39). In contrast, our findings suggest functional anatomical differences, particularly within the high-frequency gamma band (HFB), with patterns resembling those observed in fMRI research. One possible explanation for this discrepancy in results could be the use of density-based clustering analyses, which restricted the cortical tracking effect to regions with a higher concentration of significant electrodes. This approach may have revealed subtle differences in the cortical organization of the tracking effect that may have been missed without accounting for the spatial distribution of statistically significant electrodes. Alternatively, the divergence in results may stem from differences in stimulus design. The previous study used unimodal auditory stimuli, whereas our study employed audiovisual stimuli. In our study, the inclusion of multimodal stimuli may have engaged association cortices or introduced attentional competition between modalities, potentially leading to differences in how music and speech signals were integrated and processed.

### Increased cortical tracking of speech across frequency bands

The net effect of cortical tracking was higher for speech than for music. This was reflected in a higher proportion of electrodes tracking the speech envelope in both frequency bands, as well as in the results of our joint linear mixed effect models, which show a main effect of stimulus type in predicting mean cross-correlation coefficient values. Such findings are aligned with the recent literature using naturalistic stimuli, which have also shown that brain signals recover the envelope of speech signals more faithfully than that of music signals (17,35,39). Importantly, Zuk and colleagues have previously demonstrated that increased tracking of speech compared to music cannot be attributed to spectral differences in the envelope of both signals (17). Moreover, power spectra of our stimuli show that, across the different acoustic segments, speech is less rhythmic than music, showing no clear dominating peak. This suggests that increased tracking of speech cannot be attributed to higher regularity in the amplitude modulations of the speech envelope compared to the music envelope.

While our methods, as well as the lack of data about behavioral performance, prevent us from investigating the role of higher order cognitive processes, we cannot rule out the possibility that the observed effect might have a cognitive explanation. Both speech intelligibility and musical training have been shown to increase cortical tracking of the corresponding signals (13,16,22,28,86), which implies that previous experience processing a particular stimulus can modulates the magnitude of the cortical tracking effect. Additionally, attentional allocation has been associated with increased cortical tracking in previous studies (29,30,87,88).

### Cortical gradient in lag values during HFB tracking of speech signals

Our analyses also revealed interesting temporal dynamics during cortical tracking of speech stimuli in the range of HFB activity. For music and speech in the SFB, mean and median lag values were close to zero across all anatomical regions of interest, thus suggesting that neural activity in the slow oscillatory range tracks fast temporal modulations in the incoming acoustic signal in a relatively time-locked manner. Similarly, median temporal lags during HFB tracking followed the music signal by approximately 60ms on average. This could additionally reflect fast neural responses to specific acoustic features present in the music signal. In line with this, previous EcoG research has demonstrated that HFB power within prefrontal, pre/postcentral, and temporal regions encodes features such as loudness, harmonization, and the presence or absence of lyrics (78). In the same study, onset of lyrics was characterized by additional increases in HFB power, localized to posterior portions of the STG.

For speech, estimated marginal means show that HFB tracking always occurred at later lags compared to the SFB. Our results showed an increasing gradient of lag values from prefrontal, parietal and middle temporal to superior temporal regions, suggesting a highly localized effect of late-latency HFB tracking of speech within middle and posterior portions of the STG. The joint LME model further confirmed that HFB tracking of speech occurred significantly later compared to tracking of speech in the SFB, and to tracking of music in both the SFB and HFB in the three anatomical regions included in the analysis (STG, MTG and SG). This HFB gradient, which is observed for speech but not for music, could reflect tracking of amplitude modulations in the acoustic envelope at different timescales, perhaps reflecting the encoding of hierarchical structures at multiple levels. If this was the case, a prediction would be that, among trained musicians, a similar gradient in lag values across functionally relevant anatomical regions should be observed, reflecting their ability to extract structural units embedded in music signals. However, further research is needed to test this hypothesis.

### Limitations

Our study has some important limitations. First, we lack any behavioral measure that facilitates the interpretation of observed differences in cortical tracking. Without such data, we cannot rule out potential confounding factors, such as differential attentional allocation (29,30,88) due to differences in the semantic context of visual stimuli accompanying music and speech segments. Additionally, the presentation of a concurrent visual stream during passive auditory perception of music and speech segments may have introduced another confound. Previous studies have shown that rhythmic modulations in visual stimuli can crossmodally activate auditory areas (63,89,90). As a result, the effects we observed may have been influenced by the integration of visual stimuli with the auditory stimuli, potentially differing in how music and speech were processed.

Another limitation of our study is the use of a relatively broad temporal window (4000ms) for cross-correlation analyses. While this approach may have reduced statistical power and increased the likelihood of spurious results, it was motivated by two considerations. First, previous research has focused primarily on shorter temporal modulations, such as beat, notes, syllables, and words, which are often analyzed using smaller windows (around 500ms). This approach overlooks slower temporal structures such as musical and speech phrases, which occur at much slower rates. Second, a prior study using the same dataset demonstrated that longer segments (∼6 seconds) yielded better performance in predicting brain signals using Artificial Neural Networks (ANN), with improved generalization to new stimuli (54). Therefore, we chose a broader window in hopes of revealing temporal dynamics that may have been overlooked in previous studies.

Our study was also limited by the localization of the ECoG grids, which were predominantly placed in the left hemisphere. This constraint prevents us from exploring potential differences in the cortical tracking of music and speech in right hemisphere regions, which have previously been implicated in music perception and could partially explain our results. Additionally, a technical limitation involves the re-referencing of the ECoG electrodes to the average of each grid. While this is a common practice in ECoG studies, it is susceptible to field spread, meaning that activity observed in central regions could be influenced by activity from auditory areas.

We derive our interpretations from marginal mean estimates obtained from LME models. While this allows us to draw group-level conclusions, it does not capture the considerable intra-individual variability in lag estimates, which include both positive and negative lags across regions and conditions. Negative lags might reflect both higher-order and low-level predictive processing. Indeed, prior research suggests that morphosyntactic information facilitates predictive processing during speech comprehension (22,91,92), while neural mechanisms involved in rhythm perception contribute to temporal predictions during processing of music stimuli (11,93–95). Unfortunately, our methods do not allow us to determine the extent to which negative lag values reflect prediction, and any interpretation we could offer would be speculative. However, they were included in our analyses because there is enough theoretical ground to assume that they reflect relevant neurobiological and cognitive processes that support the perception of music and speech.

Relatedly, the neurobiological processes behind cortical tracking of speech and music signals remain a topic of ongoing debate. Two prominent perspectives suggest that cortical tracking of acoustic envelopes either represents the entrainment of endogenous oscillations or reflects a series of evoked responses to amplitude fluctuations within the acoustic signal (25,40,96–101). While our methods are not suited to test either of these hypotheses, our results provide evidence of differences in the temporal dynamics and cortical organization of low-level processing of acoustic envelopes, regardless of the underlying mechanism. Additionally, we do not directly address the specific information being tracked within each temporal bin and anatomical region in this work. This is important because different structural units, acoustic features, or information content can be differentially tracked by the human brain. While this question has been addressed elsewhere in the literature (38,39,49,78,86,102), we hope that the open-access nature of the data, along with improved methods to disentangle the contribution of low-level features and higher-order information to cortical tracking, will motivate others to address similar questions using this dataset.

Finally, several limitations of this study stem from the adoption of a mathematically simple approach to quantifying cortical tracking. This simplicity limits the method’s ability to disentangle the underlying neurobiological mechanisms or precisely identify the acoustic features or information tracked by neural signals, which in turn limits the depth of interpretation we can provide for our results. Nevertheless, the consistency of our findings with those of prior studies employing more sophisticated methodologies underscores the potential of our approach. Importantly, this simplicity may hold significant advantages for translational applications, particularly in contexts where minimizing computational demands is a priority or where theoretical assumptions are less relevant for practical applications.

## Conclusion

In summary, our results show widespread cortical tracking and reveal both functional overlap and anatomical specialization in the spatial and temporal dynamics associated with the tracking of naturalistic music and speech stimuli. Passive perception of music and speech was associated with distributed tracking of acoustic signals in the SFB with near zero delays in mostly overlapping temporal and perisylvian regions. While some overlap was observed in the HFB for both stimulus types, marked anatomical and functional selectivity emerged, with regions such as the dlPFC and IPL areas preferentially tracking music, and the IFG preferentially tracking speech. Notably, the overall magnitude of the tracking effect was higher for speech across both frequency bands. Additionally, a gradient in lag values was observed during HFB tracking of speech, spanning from association areas in prefrontal, middle temporal, and parietal regions towards STG areas, which was nonetheless absent during HFB tracking of music.

Our findings further support previous research indicating that cortical tracking of music and speech extends beyond controlled experimental settings to more complex naturalistic signals, which are far less controlled, less rhythmic, and that are presented under conditions of sensory multimodality. Moreover, our results highlight the brain’s use of both domain-general mechanisms and frequency-dependent anatomical specialization when tracking natural acoustic signals. Finally, we highlight a complex interaction between stimulus type, cortical region, and frequency band, which could underlie the brain’s ability to extract hierarchical structures from the envelope of natural acoustic signals. However, future research is needed to empirically test this hypothesis.

## Data availability statement

Data for this study is open and can be accessed at https://openneuro.org/datasets/ds003688. Code used for data analysis can be made available upon request.

