## Supplementary Material for "Anatomically distinct cortical tracking of music and speech by slow (1-8Hz) and fast (70-120Hz) oscillatory activity"

**This PDF includes:**

Supplementary tables s1, s2, s3, s4, s5

Supplementary figures s1, s2 s3, s4, s5, s6

| Subject | Sex | Age | Handedness | Language Dominance | Language dominance technique | Grid hemisphere | HD Grid |
|---|---|---|---|---|---|---|---|
| sub-02 | F | 9 | R | L | fTCD + fMRI | L | no |
| sub-03 | F | 33 | R | L | Wada | L | no |
| sub-05 | F | 33 | R | L | fTCD | L | no |
| sub-06 | F | 43 | R | L | fMRI | L | no |
| sub-10 | M | 8 | R | L | Wada | L | no |
| sub-12 | M | 37 | R | L | ECS | L | no |
| sub-16 | M | 17 | R | L | Wada | L | no |
| sub-18 | F | 15 | R | L | fMRI | L | no |
| sub-19 | F | 8 | R | possibly L | fMRI | L | no |
| sub-20 | F | 25 | L | L | Wada | L | no |
| sub-22 | M | 21 | R | L | fMRI | L | no |
| sub-24 | F | 47 | R | L | Wada | L | no |
| sub-25 | M | 14 | R | L | fTCD | L | no |
| sub-26 | F | 48 | R | L | fMRI | L | no |
| sub-27 | M | 15 | R | L | fMRI | L | no |
| sub-34 | F | 51 | R | L | ECS | L | no |
| sub-36 | F | 52 | R | L | fMRI | L | yes |
| sub-39 | F | 5 | R | possibly L | ECS (fTCD inconclusive, maybe bilateral) | L | no |
| sub-40 | M | 49 | R | L | fMRI | L | no |
| sub-45 | M | 19 | R | L | fMRI | L | yes |
| sub-46 | F | 41 | L | L | fMRI | L | no |
| sub-48 | F | 18 | R | L | fTCD and/or fMRI | L | no |
| sub-51 | M | 46 | R | L | fMRI + fTCD | L | no |
| sub-54 | F | 31 | R | L | fMRI | L | no |
| sub-55 | F | 23 | R | L | Wada | L | no |
| sub-58 | F | 16 | R | L | fMRI | L | no |
| sub-59 | F | 30 | L | L | fMRI | L | no |
| sub-60 | M | 42 | R | L | Wada | L | yes |
| sub-61 | F | 16 | R | L | fTCD | L | no |
| sub-63 | M | 5 | n/a | L | ECS | L | no |

**Table s1.** Summary of demographic and clinical information for the subjects included in this study. For reproducibility, subject identifiers correspond to those used in the original dataset.

| Electrodes per subject | | | |
|---|---|---|---|
| | *Total* | *After statistics* | *Percentage* |
| sub-02 | 47 | 15 | 31.91% |
| sub-03 | 80 | 37 | 46.25% |
| sub-05 | 60 | 28 | 46.67% |
| sub-06 | 63 | 26 | 41.27% |
| sub-10 | 48 | 29 | 60.42% |
| sub-12 | 64 | 29 | 45.31% |
| sub-16 | 93 | 59 | 63.44% |
| sub-18 | 60 | 31 | 51.67% |
| sub-19 | 69 | 46 | 66.67% |
| sub-20 | 54 | 27 | 50.00% |
| sub-22 | 45 | 23 | 51.11% |
| sub-24 | 60 | 33 | 55.00% |
| sub-25 | 48 | 26 | 54.17% |
| sub-26 | 47 | 23 | 48.94% |
| sub-27 | 62 | 28 | 45.16% |
| sub-34 | 60 | 30 | 50.00% |
| sub-36 | 121 | 43 | 35.54% |
| sub-39 | 64 | 36 | 56.25% |
| sub-40 | 47 | 37 | 78.72% |
| sub-45 | 107 | 62 | 57.94% |
| sub-46 | 41 | 16 | 39.02% |
| sub-48 | 61 | 36 | 59.02% |
| sub-51 | 44 | 20 | 45.45% |
| sub-54 | 52 | 43 | 82.69% |
| sub-55 | 72 | 55 | 76.39% |
| sub-58 | 80 | 50 | 62.50% |
| sub-59 | 40 | 21 | 52.50% |
| sub-60 | 73 | 23 | 31.51% |
| sub-61 | 48 | 30 | 62.50% |
| sub-63 | 48 | 34 | 70.83% |
| *Sum* | 1858 | 996 | 53.61% |
| *Mean* | 61.93 | 33.20 | |
| *SD* | 19.09 | 12.00 | |

**Table s2.** Total number of electrodes per subject before and after permutation statistics, and percentage of statistically significant electrodes for each subject.

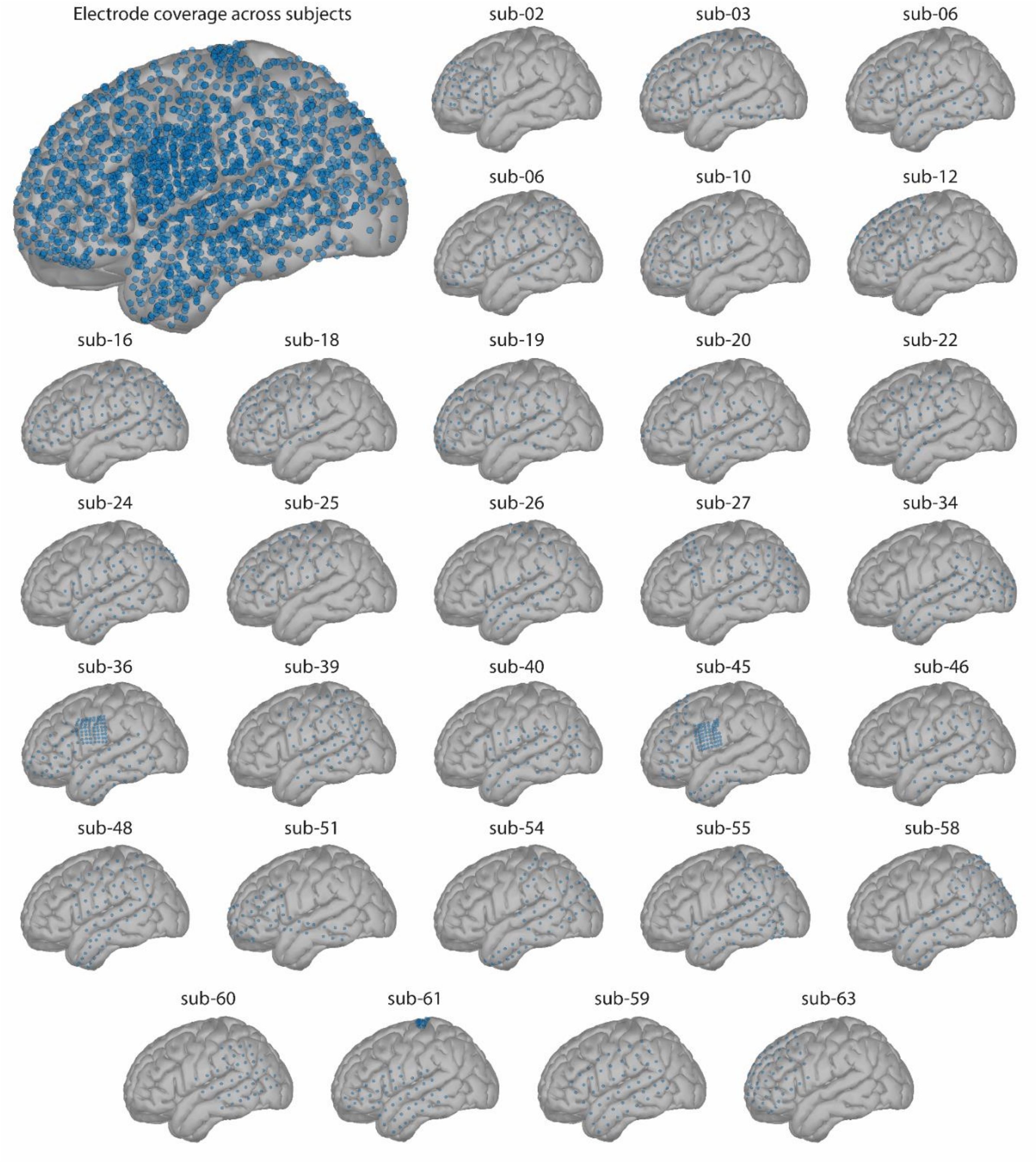


**Figure s1.** Electrode coverage across all subjects (n = 30, top-left panel) and electrode location for each individual subject included in the analyses, plotted against the MNI ICBM152 cortical template.

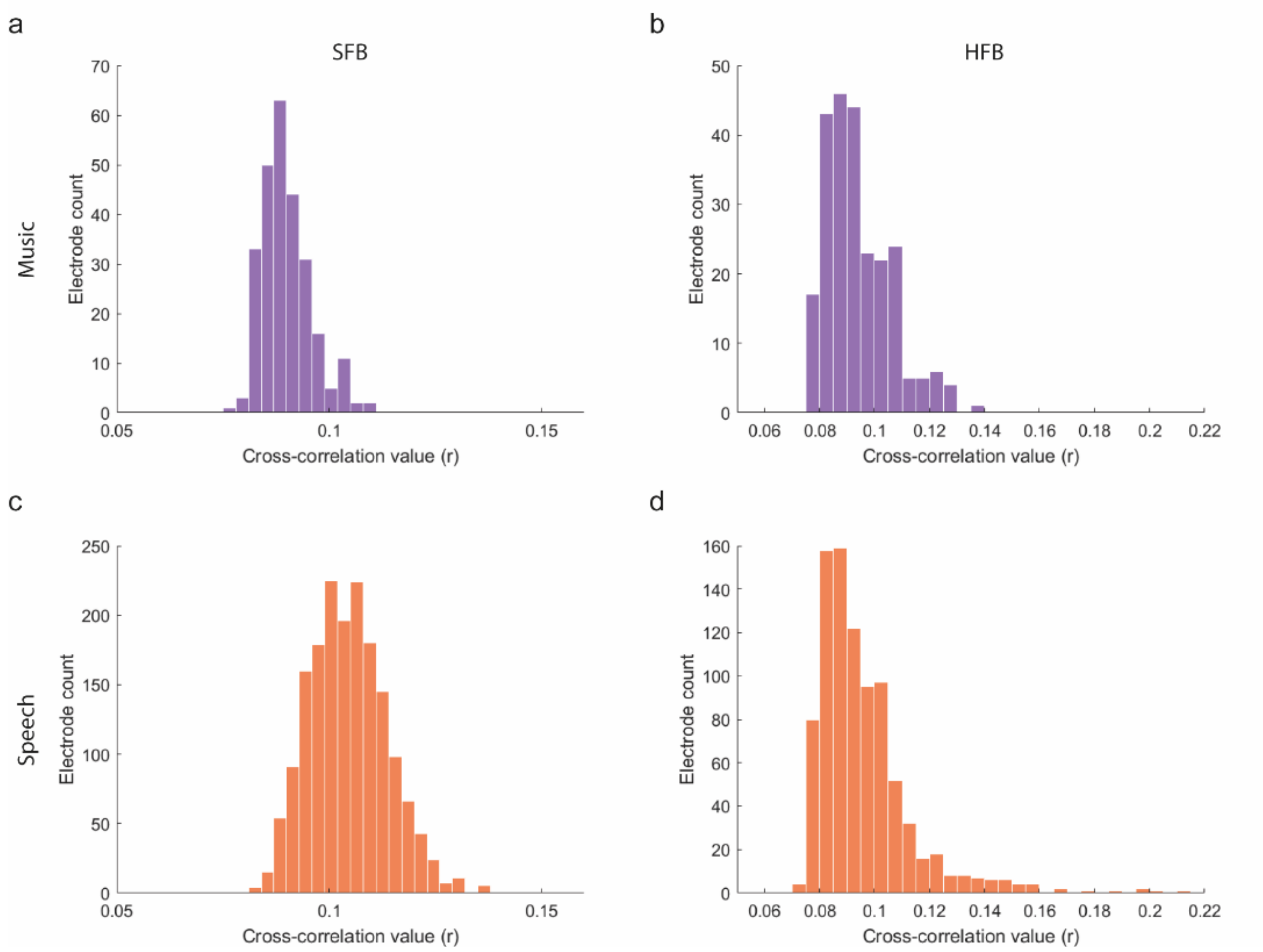


**Figure s2.** Distribution of observed cross-correlation coefficients (pooled for all participants) prior to statistics for music (purple) and speech (orange) in the SFB (a, c) and HFB (b,d) frequency ranges.

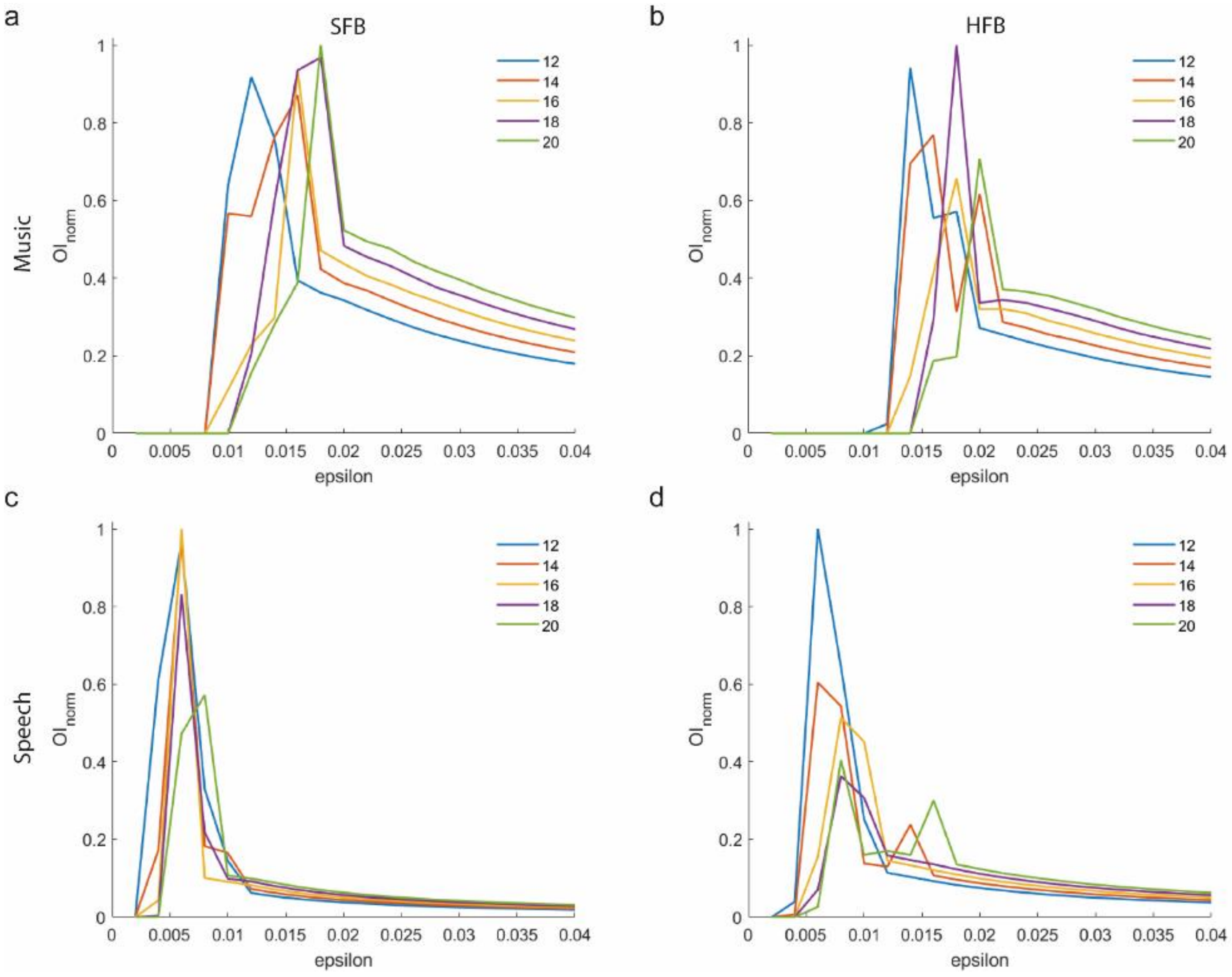


**Figure s3.** Optimization index as a function of parameter combination in the DBSCAN analysis. X axis shows different epsilon parameters and different colored lines illustrate different minimum points. The optimization procedure was conducted separately for each frequency band and condition. Optimal parameters maximize the number of electrodes and minimizes epsilon.

| **Model** | **Response variable** | | | |
|---|---|---|---|---|
| | *r* | | *Lag* | |
| | *AIC* | *BIC* | *AIC* | *BIC* |
| (1\|subject) | -1293.76 | -1283.81 | 277.78 | 287.73 |
| frequency + (1\|subject) | -1299.13 | -1285.85 | 279.29 | 292.57 |
| region + (1\|subject) | -1291.16 | -1271.26 | 279.31 | 299.22 |
| frequency + region + (1\|subject) | -1299.08 | -1275.85 | 279.35 | 302.58 |
| frequency * region + (1\|subject) | -1293.21 | -1260.03 | 280.13 | 313.31 |
| frequency * region + (1\|subject:region) | -1294.50 | -1261.32 | 279.93 | 313.11 |
| frequency * region + (1\|subject) + (1\|subject:region) | -1292.88 | -1256.38 | 281.79 | 318.28 |

**Table s3.** Performance of LME analysis of music for the two response variables of interest (correlation coefficient and temporal lag). For cross-correlation coefficients during cortical tracking of music, a single predictor LME model including frequency as fixed effect and random intercepts for subject outperformed the null model ($X^2(1) = 7.36$, p = 0.0007), but the single predictor model for cortical regions did not. Adding both predictors as fixed effects in the same model did not improve model fit against the frequency-only model, nor did adding an interaction term between them. Finally, adding cortical region and the subject-by-region interaction did not improve model fit either. For temporal lags, no model was significantly better than the intercept-only model.

| Model | Response variable | | | |
|---|---|---|---|---|
| | *r* | | *Lag* | |
| | *AIC* | *BIC* | *AIC* | *BIC* |
| (1\|subject) | -6261.43 | -6246.38 | 1805.96 | 1821.01 |
| frequency + (1\|subject) | -6264.92 | -6244.85 | 1644.32 | 1664.39 |
| region + (1\|subject) | -6298.25 | -6258.12 | 1790.98 | 1831.11 |
| frequency + region + (1\|subject) | -6309.98 | -6264.83 | 1640.41 | 1685.55 |
| frequency * region + (1\|subject) | -6333.81 | -6263.57 | 1617.03 | 1687.26 |
| frequency * region + (1\|subject:region) | -6349.42 | -6279.19 | 1617.92 | 1688.15 |
| frequency * region + (1\|subject) + (1\|subject:region) | -6351.47 | -6276.22 | 1611.49 | 1686.74 |

**Table s4.** Performance of LME analysis of speech for the two response variables of interest (correlation coefficient and temporal lag). For cross-correlation values during cortical tracking of speech, the single predictor models for frequency ($X^2(1) = 5.48$, p = 0.019) and region ($X^2(5) = 46.82$, p = 6.18e-09) outperformed the null model. Adding the interaction between both predictors further improved fit against the single-predictor model for region ($X^2(6) = 47.55$, p = 1.45e-08) and an additive model including both predictors ($X^2(5) = 33.82$, p = 2.57e-06). Finally, adding subject-by-region interaction to random intercept for subject as random effects further improved the interaction model fit ($X^2(1) = 19.66$, p = 9.242-06). For temporal lags, both single predictor models for frequency ($X^2(1) = 163.64$, p = 2.2e-16) and region ($X^2(5) = 24.98$, p = 0.0001) outperformed the null model. Adding both predictors within the same model further improved model fit compared to the frequency ($X^2(5) = 13.19$, p = 0.016) and region ($X^2(5) = 152.57$, p = 2.2e-16) single-predictor models, and adding the interaction term significantly improved fit compared to the additive model ($X^2(5) = 33.74$, p = 3.17e-06). Finally, an additional random effect for the interaction of subject-by-region further improved model fit ($X^2(1) = 7.54$, p = 0.006).

| Model | Response variable | | | |
|---|---|---|---|---|
| | *r* | | *Lag* | |
| | *AIC* | *BIC* | *AIC* | *BIC* |
| (1\|subject) | -4371.14 | -4357.07 | 1345.94 | 1360.01 |
| frequency + (1\|subject) | -4369.15 | -4350.39 | 1245.38 | 1264.14 |
| region + (1\|subject) | -4386.77 | -4363.32 | 1338.46 | 1361.90 |
| condition + (1\|subject) | -4479.78 | -4461.02 | 1344.27 | 1363.02 |
| frequency + region + condition + (1\|subject) | -4490.98 | -4458.16 | 1236.00 | 1268.82 |
| frequency * region * condition + (1\|subject) | -4492.39 | -4426.73 | 1191.97 | 1257.62 |
| frequency * region * condition + (1\|subject:region) | -4501.54 | -4435.89 | 1190.03 | 1255.68 |
| frequency * region * condition + (1\|subject) + (1\|subject:region) | -4502.26 | -4431.91 | 1191.89 | 1262.23 |

**Table s5.** Performance of LME joinst (music and speech) model for the two response variables of interest (correlation coefficient and temporal lag). A joint LME analysis was conducted for both music and speech including the cortical regions where electrodes were found across these conditions. In this model, stimulus type, frequency band and cortical regions were fixed effects of interest, and the subject, region and subject-by-region interaction were random effects of interest. We conducted forward stepwise model selection procedure as reported above, using the same response variables of interest. For cross-correlation coefficients, the single predictor models for region ($X^2(2) = 19.63$, p = 5.45e-05) and condition ($X^2(1) = 110.64$, p = 2.2e-16) showed a significantly better fit compared to the null model. Adding the interaction term between the three predictors improved all single-predictor models (frequency: $X^2(10) = 143.24$, p = 2.2e-16; region: $X^2(9) = 123.61$, p = 2.2e-16, condition: $X^2(10) = 35.61$, p = 0.0003) as well as the additive model ($X^2(7) = 14.40$, p = 0.031). Adding the cortical region as random intercept on top of subject did not improve model fit, but adding the region-by-subject interaction did ($X^2(1) = 22.87$, p = 0.0006). For temporal lags, single predictor models for frequency ($X^2(1) = 102.56$, p =2.2e-16) and region ($X^2(2) = 11.49$, p = 0.0032) showed better fit compared to the null model, but not the single-predictor model for condition did not. Adding the three predictors as fixed effect in the same model further improved fit compared to all these models and adding the interaction term between them further improved fit compared to the additive model ($X^2(7) = 58.03$, p = 3.72e-10). Adding the region or the subject-by-region interaction as random effects did not improve fit of the interaction model.

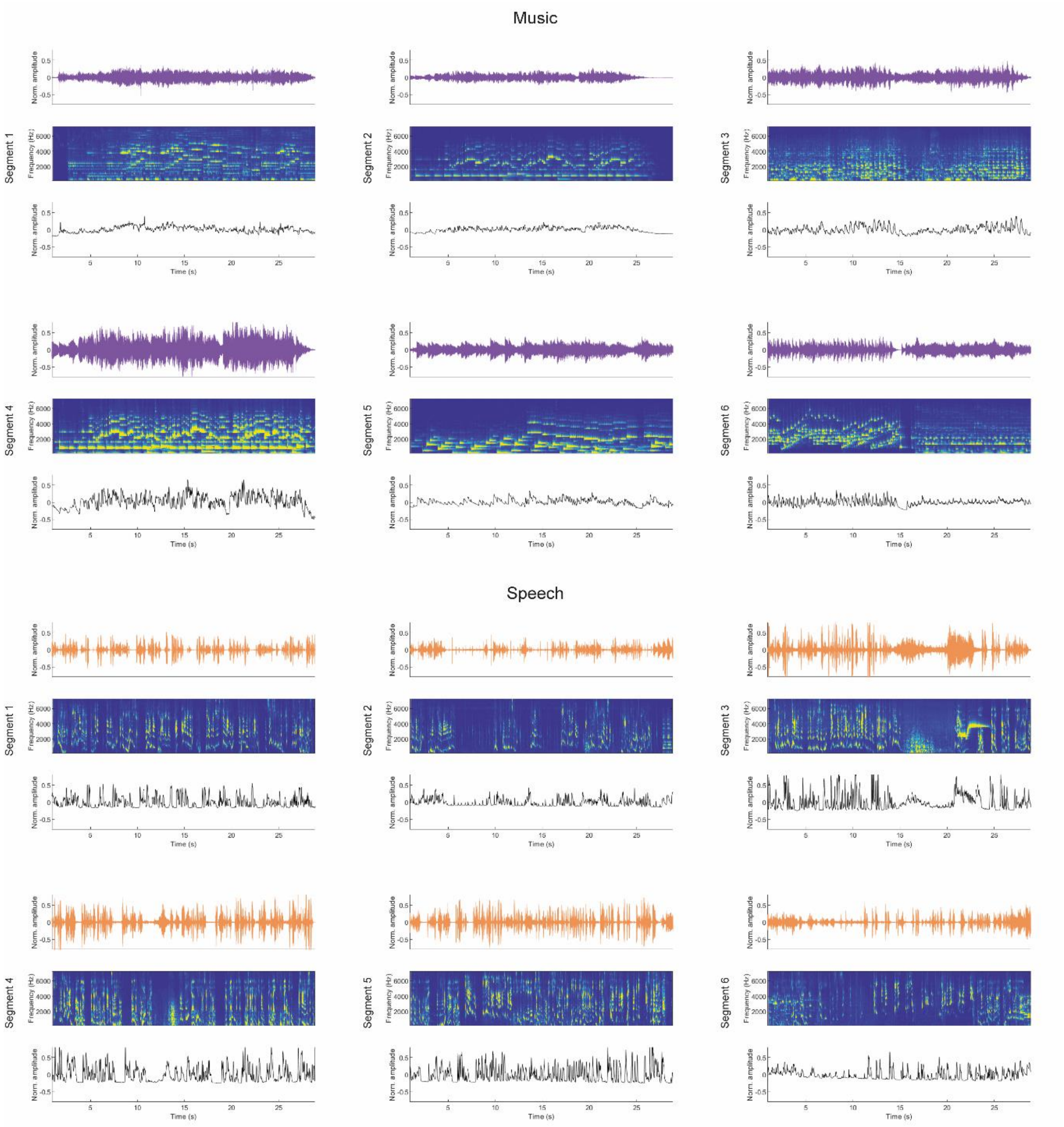


**Figure s4.** Soundwaves (top), spectrogram (middle) and envelope (bottom) of the six music and speech segments used as stimuli.

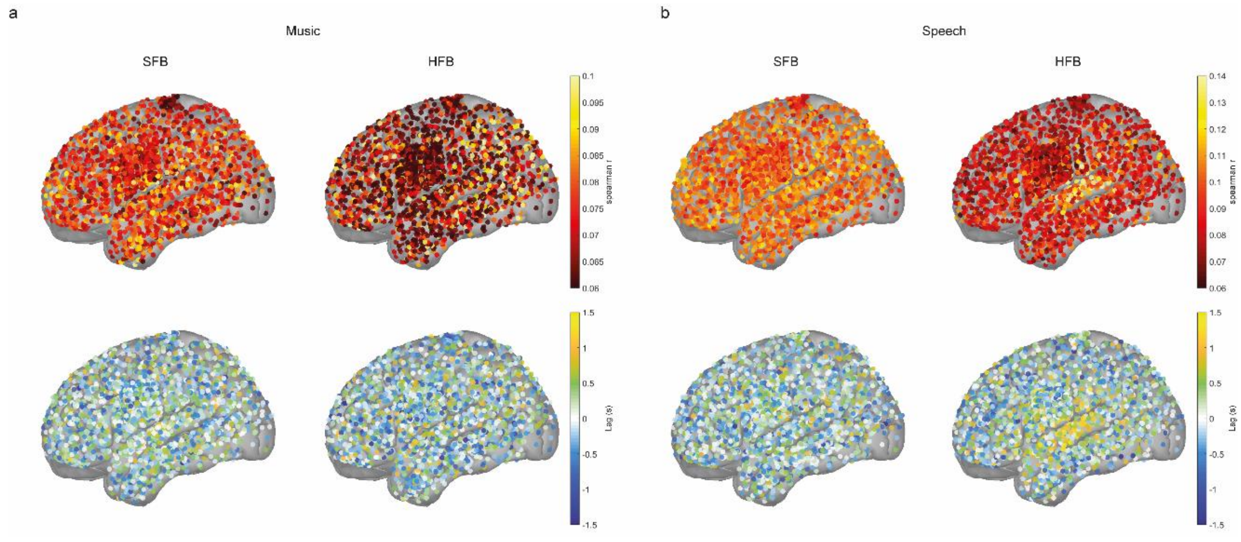


**Figure s5.** Non-thresholded estimates of cross-correlation coefficients (top row) and temporal lags (bottom row) for music (**a**) and speech (**b**) in both frequency bands of interest. This figure illustrates how our statistical analyses affect the results reported in our work.

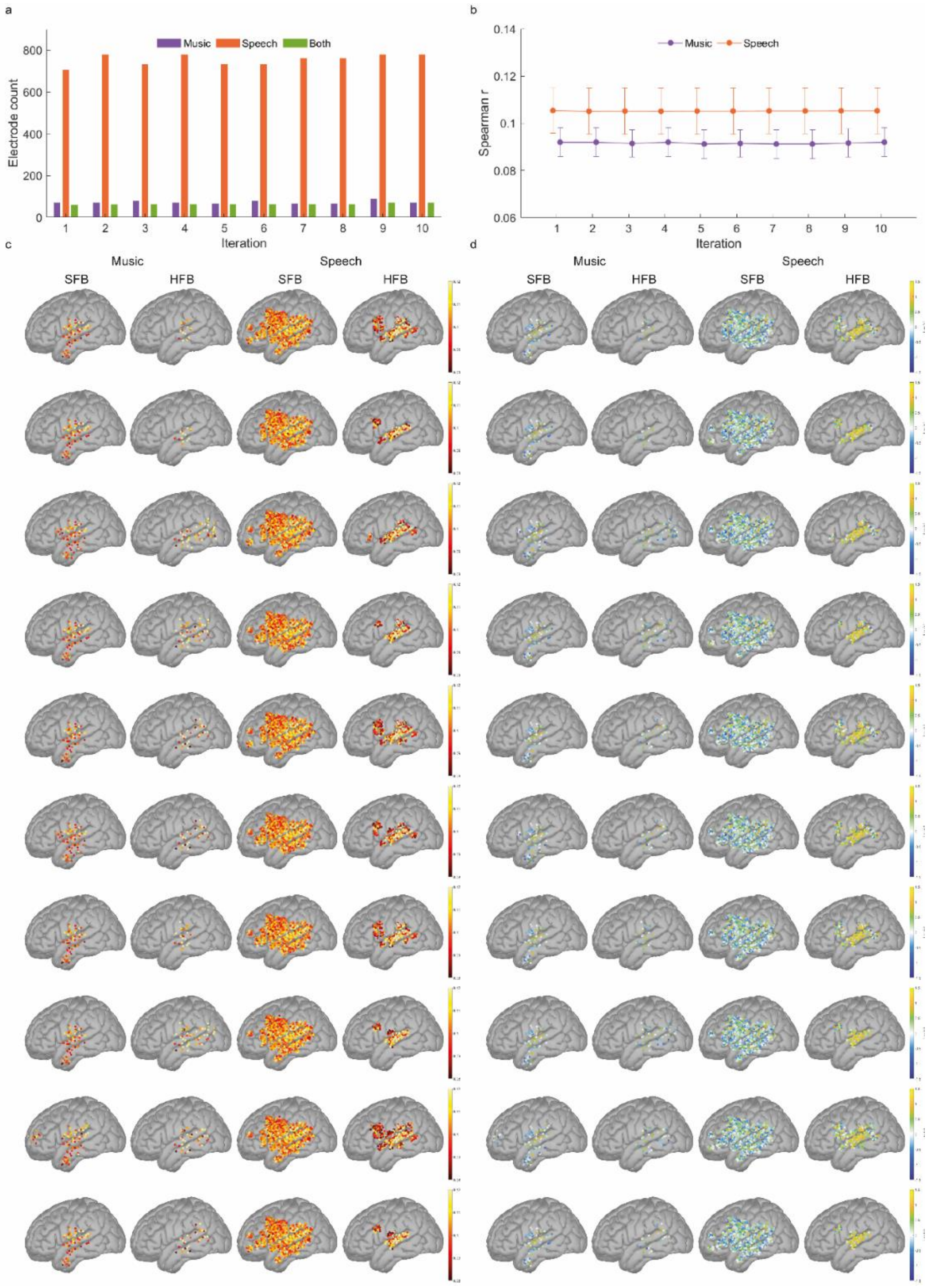


**Figure s6**. To investigate the consistency of our results across stimulus type, data was reanalyzed over 10 iterations using the same procedures as reported in the main set of analyses. For each iteration, we randomly excluded two segments per condition, corresponding to 33% of the data. **a.** Shows the number of statistically significant electrodes in each iteration. **b.** Magnitude of the effect in each iteration. **c.** and **d.** Cortical organization of the magnitude of the effect (c) and temporal lags (d). Each row corresponds to a different iteration.
